# AUTOPILOT: *Automating experiments with lots of Raspberry Pis*

**DOI:** 10.1101/807693

**Authors:** Jonny L. Saunders, Lucas A. Ott, Michael Wehr

**Affiliations:** University of Oregon, Institute of Neuroscience, Department of Psychology, Eugene, OR 97403, United States

## Abstract

Neuroscience needs behavior, and behavioral experiments require the coordination of large numbers of heterogeneous hardware components and data streams. Currently available tools strongly limit the complexity and reproducibility of experiments. Here we introduce Autopilot, a complete, open-source Python framework for experimental automation that distributes experiments over networked swarms of Raspberry Pis. Autopilot enables qualitatively greater experimental flexibility by allowing arbitrary numbers of hardware components to be combined in arbitrary experimental designs. Research is made reproducible by documenting all data and task design parameters in a human-readable and publishable format at the time of collection. Autopilot provides a high-level set of programming tools while maintaining submillisecond performance at a fraction of the cost of traditional tools. Taking seriously the social nature of code, we scaffold shared knowledge and practice with a publicly editable semantic wiki and a permissive plugin system. Autopilot’s flexible, scalable architecture allows neuroscientists to work together to design the next generation of experiments to investigate the behaving brain.

## 1 Introduction

Animal behavior experiments need precision and patience, so we make computers do them for us. The complexity of contemporary behavioral experiments, however, presents a stiff methodological challenge. For example, researchers might wish to measure pupil dilation[2], respiration[3], and running speed[4], while tracking the positions of body parts in 3 dimensions[5] and recording the activity of large ensembles of neurons[6], as subjects perform tasks with custom input devices such as a steering wheel[7] while immersed in virtual reality environments using stimuli synthesized in real time[8, 9]. Coordinating the array of necessary hardware into a coherent experimental design—with the millisecond precision required to study the brain—can be daunting.

Historically, researchers have developed software to automate behavior experiments as-needed within their lab or relied on purchasing proprietary software (eg. [10]). Open-source alternatives have emerged recently, often developed in tandem with hardware peripherals available for purchase [11, 12]. However, the diverse hardware and software requirements for behavioral experiments often lead researchers to cobble together multiple tools to perform even moderately complex experiments. Understandably, most software packages do not attempt to simultaneously support custom hardware operation, behavioral task logic, stimulus generation, and data acquisition. The difficulty of designing and maintaining lab-idiosyncratic systems thus defines much of the everyday practice of science. Idiosyncratic systems can hinder reproducibility, especially if the level of detail reported in a methods section is sparse[13]. Additionally, development time and proprietary software are expensive, as are the custom hardware peripherals that are required to use most available opensource behavior software, stratifying access to state-of-the-art techniques according to inequitable funding distributions.

Technical challenges are never merely technical: they reflect and are structured by underlying *social* challenges in the organization of scientific labor and knowledge work. Lab infrastructure occupies a space between technology intended for individual users and for large organizations: that of *groupware*^1^ [15, 14]. Experimental frameworks thus face the joint challenge of technical competency while also embedding in and supporting existing cultures of practice. Behind every line of code is an unwritten wealth of technical knowledge needed to make use of it, as well as an unspoken set of beliefs about how it is to be used — labs aren’t born fresh on release day ready to retool at a moment’s notice, they’re held together by decades of duct tape and run on ritual. The boundaries of this “contextual knowledge” extend fluidly beyond individual labs, structuring disciplinary, status, and role systems in scientific work[16]. Given their position at the intersection of scientific theory, technical work, data production, and social organization, experimental frameworks are an elusive design challenge, but also an underexplored means of realizing some of our loftier dreams of open, accessible, and collaborative science.

Here we present Autopilot, a complete open-source software and hardware framework for behavioral experiments. We leverage the power of distributed computing using the surprisingly capable Raspberry Pi 4^2^ to allow researchers to coordinate arbitrary numbers of heterogeneous hardware components in arbitrary experimental designs.

Autopilot takes a different approach than existing systems to overcome the technical challenges of behavioral research: *just use more computers*. Specifically, the advent of inexpensive single-board computers (ie. the Raspberry Pi) that are powerful enough to run a full Linux operating system allows a unified platform to run on every Pi or other computer in the system so that they can work together seamlessly. At the core of its architecture are networking classes (Section 3.8) that are fast enough to stream electrophysiological or imaging data and flexible enough to make the mutual coordination of hardware straightforward.

This distributed design also makes Autopilot extremely scalable, as the Raspberry Pi’s $35-$75 price tag makes it an order of magnitude less costly than comparable systems (Section 2.3). Its low cost doesn’t come at the expense of performance or useability: Autopilot provides an approachable, high-level set of tools that still have input and output precision between dozens of microseconds to a few milliseconds (Sections 2.1 and 4).

Autopilot balances experimental flexibility with support. Its task design infrastructure is flexible enough to perform arbitrary experiments, but also provides support for data management, plotting task progress, and custom training regimens. We try to bridge multiple modalities of use: use its modular framework of tools out of the box, or use its complete low-level API documentation^3^ to hack it to do what you need. Rather than relying on costly proprietary hardware modules, users can take advantage of the wide array of peripherals and extensive community support available for the Raspberry Pi. Autopilot is designed to be *permissive*: build your whole experiment with it or just use its networking modules, adapt it to existing hardware, integrate your favorite analysis tool. We designed Autopilot to *play nice* with other software libraries and existing practices rather than force you to retool your lab around it.

Finally, we have designed Autopilot to help scientists do reproducible research and be good stewards of the human knowledge project. Experiments are not written as scripts that are reliant on the particularities of each researcher’s hardware configuration. Instead, we have designed the system to encourage users to write reusable, portable experiments that can be incorporated into a public central library while also allowing space to iterate and refine without needing to learn complicated programming best-practices to contribute. Every parameter that defines an experiment is automatically saved in publication-ready format, removing ambiguity in reported methods and facilitating exact replication with a single file. Its plugin system is built atop a densely-linked [semantic wiki](https://wiki-auto-pi-lot.com/)^4^ that fluidly combines human- and computer-readable, communally editable technical knowledge that surrounds your experiments with the software that performs them.

We begin by defining the requirements of a complete behavioral system and evaluating two current examples (Sections 1.1 and 1.2). We then describe Autopilot’s design principles (Section 2) and how they are implemented in the program’s structure (Section 3). We close with a demonstration of its current capabilities and our plans to expand them (Sections 4 and 5).

### 1.1 Existing Systems for Behavioral Experiments

At minimum, a complete system to automate behavioral experiments has 6 requirements:

1. Hardware to interact with the experimental subject, including **sensors** (eg. photodiodes, cameras, rotary encoders) to receive input and **actuators** (eg. lights, motors, solenoids) to provide feedback.
2. Some capability to synthesize and present sensory stimuli. Ideally both discrete stimuli, like individual tone pips or grating patches, and continuous stimuli, like those used in virtual reality experiments, should be possible.
3. A framework to coordinate hardware and stimuli as a task. Task definition should be flexible such that it facilitates rather than constrains experimental design.
4. A data management system that allows fine control of data collection and format. Data should be human readable and include complete metadata that allows independent analysis and reproduction. Ideally the program would also allow some means of realtime data processing of sensor values for use in a task.
5. Some means of visualizing data as it is collected in order to observe task status. It should be possible to customize visualization to the needs and structure of the task.
6. Finally, a user interface to control task operation. The UI should make it possible for someone who does not program to operate the system.

We will briefly describe two other systems that meet this definition of completeness: pyControl and Bpod.

#### pyControl

[pyControl](https://pycontrol.readthedocs.io/en/latest/)[17] is a behavioral framework built in Python by the Champalimaud Foundation. It uses the [micropython microcontroller](https://micropython.org/) (“pyboard”) as its primary hardware device along with several extension boards [sold by openephys](http://www.open-ephys.org/store/pycontrol). The pyboard has four I/O ports, or eight with a multiplexing expander board. Schematics are available for many other hardware components like solenoid valve drivers and rotary encoders. Multiple pyboards can be connected to a computer via USB and run independent tasks simultaneously with a GUI.

There is limited support for some parametrically defined sound stimuli, presented from a separate amplifier connected using the I2C protocol. Visual stimuli are unsupported.

Like most behavioral software, pyControl uses a finite-state machine formalism to define its tasks. A task is a set of discrete states, each of which has a set of events that transition the task from one state to another. pyControl also allows timed transitions between states, and one function that is called on every event for a rough sort of parallelism. pyControl also allows the use of external variables to control state logic, making these state machines more flexible than strict finite state machines.

All events and states are stored alongside timestamps as a plain text log file, one file per subject per session (Figure 1.1). Analog data are stored in a custom binary serialization that alternates 4-byte data and timestamp integers.

**Figure 1.1:**
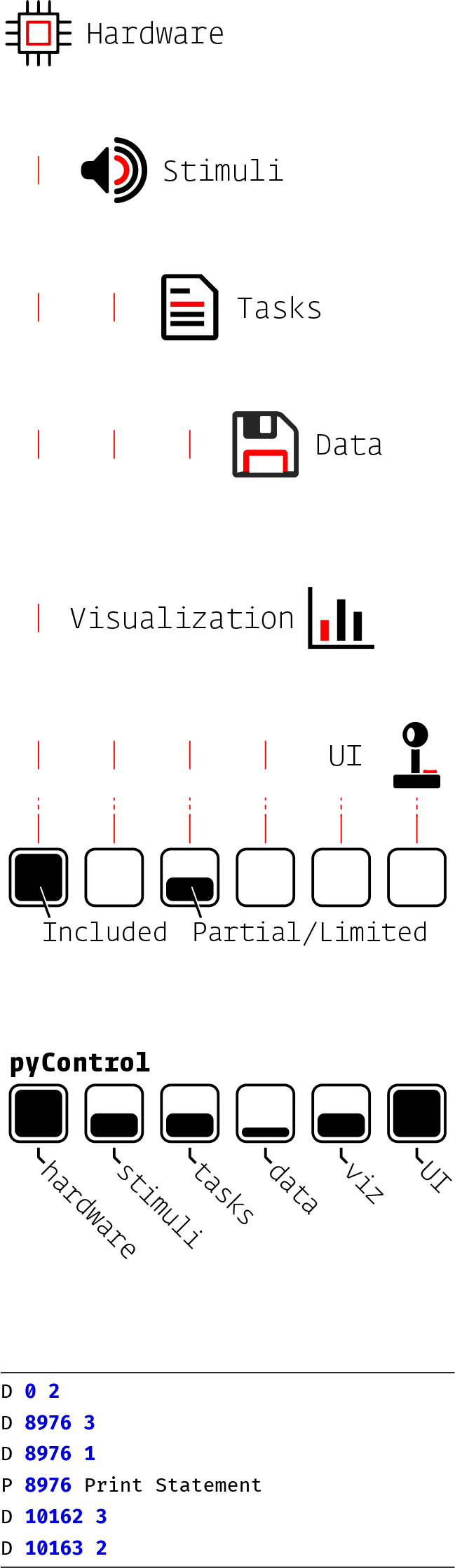
pyControl data is stored as plain text, each line having a type (like **D**ata or **P**rint), timestamp, and state

There is only one plot type available in the GUI, a raster plot of events, and no facility for varying the plot by task type. The GUI is otherwise quite capable, including the ability to batch run subjects, redefine task variables, and configure hardware.

#### Bpod

Bpod is primarily a collection of hardware designs and an assembly service run by [Sanworks LLC](https://www.sanworks.io/about/about.php). Similar to pyControl, each Bpod behavior box is based on a finite-state machine microcontroller with four I/O ports. Additional hardware modules provide extended functionality. Bpod is controlled using its own [MATLAB package](https://github.com/sanworks/Bpod_Gen2), though there are at least two other third-party software packages, [BControl](https://brodylabwiki.princeton.edu/bcontrol/index.php/Main_Page) and [pyBpod](https://pybpod.github.io/), that can control Bpod hardware. A task is implemented as a MATLAB script that constructs a new state machine for each trial, uploads it to the Bpod, and waits for the trial to finish. As a result, only one Bpod can be used per host computer, or at least per MATLAB session. Data are stored as trial-split events in a MATLAB structure.

There are a few basic plots for two-alternative forced choice tasks, but there doesn’t seem to be a prescribed way to add additional plots. Bpod has a reasonably complete GUI for managing the hardware and running tasks, but it is relatively technical (Figure 1.2).

**Figure 1.2:**
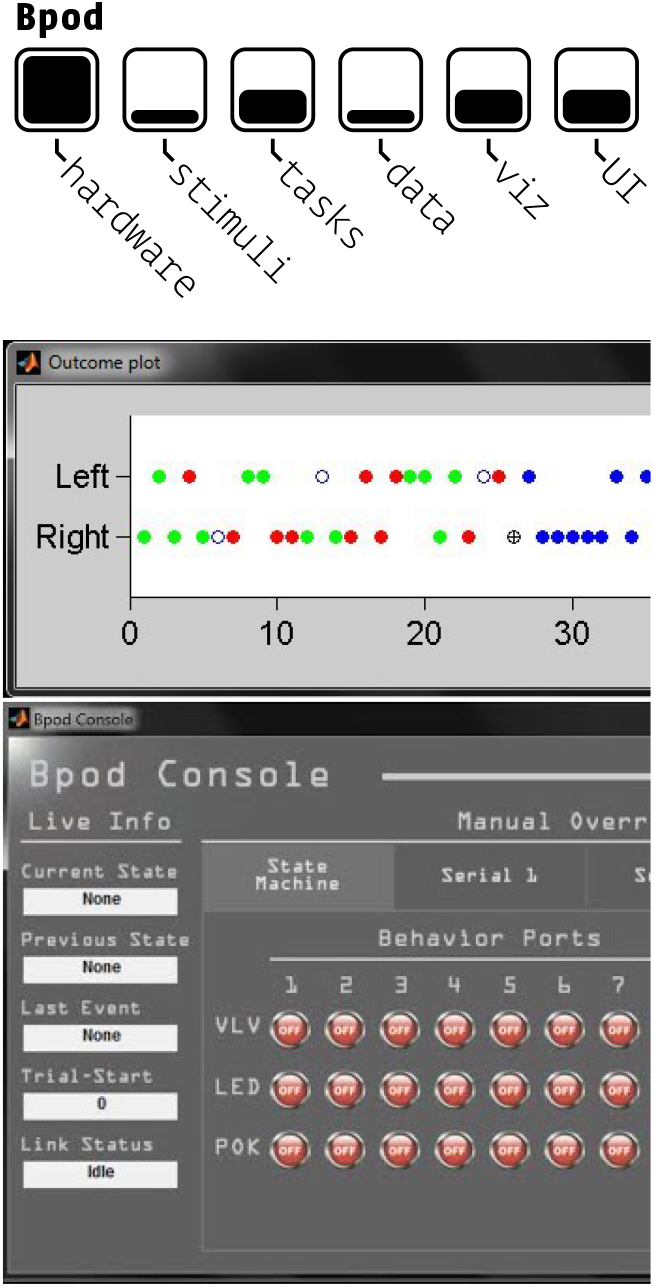
A Bpod event plot (above) showing the results of individual behavioral trials, and the Bpod GUI (below).

For brevity we have omitted many other excellent tools that perform some subset of the operations of a complete behavioral system, or otherwise have a substantial difference in scope.^5^

### 1.2 Limitations of Existing Systems

We see several limitations with these and other behavioral systems:

- **Hardware** - Both Pycontrol and Bpod strongly encourage users to purchase a limited set of hardware modules and add-ons from their particular hardware ecosystem. If a required part is not available for purchase, neither system provides a clear means of interacting with custom hardware aside from typical digital inputs and outputs, requiring the user to ‘tack on’ loosely-integrated components. There is also a hard limit on the *number* of hardware peripherals that can be used in any given task, as there is no ability to use additional pyboards or Bpod state machines in a single task. The microcontrollers used in these systems also impose strong limits on their software: neither run a full, high-level programming language^6^. We will discuss this further in section 2.2. A broader limitation of existing systems is the difficulty of flexibly integrating diverse hardware with the analytical tools necessary to perform the next generation of behavioral neuroscience experiments that study “naturalistic, unrestrained, and minimally shaped behavior”[25].
- **Stimuli** - Stimuli are not tightly integrated into either of these systems, requiring the user to write custom routines for their synthesis, presentation, and description in the resulting data. Neither are capable of delivering visual stimuli. Since the publication of the initial version of this manuscript, Bpod has added support for a HiFiberry sound card that we also describe here[26], but the sound generation API appears to be [unchanged](https://github.com/sanworks/Bpod_Gen2/blob/df6cd0c7d5df8247b02077b05fc263f79b86b096/Examples/Protocols/Sound/HiFiSound2AFC_TrialManager/HiFiSound2AFC_TrialManager.m), with a single method for generating [sine waves](https://github.com/sanworks/Bpod_Gen2/blob/1cb181dffbb7394acd18819f1d268fd9dec6ec5b/Functions/Internal%20Functions/GenerateSineWave.m). Some parametric audio stimuli are included in the [pyControl source code](https://github.com/pyControl/code/blob/master/pyControl/audio.py) but we were unable to find any documentation or examples of their use.
- **Tasks** - Tasks in both systems require a large amount of code and effort duplication. Neither system has a notion of reusable tasks or task ‘templates,’ so every user typically needs to rewrite every task from scratch. Bpod’s structure in particular tends to encourage users to write long [task scripts](https://github.com/sanworks/Bpod_Gen2/blob/df6cd0c7d5df8247b02077b05fc263f79b86b096/Examples/Protocols/Sound/AnalogSound2AFC/AnalogSound2AFC.m) that contain the entire logic of the task including updating plots and recreating state machines (Figure 1.3). Since there is little notion of how to share and reuse common operations, most users end up creating their own secondary libraries and writing them from scratch. Another factor that contributes to the difficulty of task design in these systems is the need to work around the limitations of finite state machines, which we discuss further in section 3.3.
- **Data** - Data storage and formatting is basic, requiring extensive additional processing to make it human readable. For example, to determine whether a subject got a trial correct in an [example](https://github.com/sanworks/Bpod_Gen2/blob/master/Examples/Protocols/Light/Light2AFC/Light2AFC.m) Bpod experiment, one would use the following code:

~~~
SessionData.RawEvents.Trial{**1**, **1**}.States.Punish(**1**) ~= NaN
~~~ As a result, data format is idiosyncratic to each user, making data sharing dependent on manual annotation and metadata curation from investigators.
- **Visualization & GUI** - The GUIs of each of these systems are highly technical, and are not designed to be easily used by non-programmers, though pyControl’s documentation offsets much of this difficulty. Visualization of task progress is quite rigid in both systems, either a timeseries of task states or plots specific to two-alternative forced choice tasks. In the examples we have seen, adapting plots to specific tasks is mostly ad-hoc use of external tools.
- **Documentation** - Writing good documentation is challenging, but particularly for infrastructural systems where a user is likely to need to modify it to suit their needs it is important that it be possible to understand its lower-level workings. PyControl has relatively good [user documentation](https://pycontrol.readthedocs.io/en/latest/) for how to use the system, but no API-level documentation. Bpod’s [documentation](https://sites.google.com/site/bpoddocumentation/home) is a bit more scattered, and though it does have documentation for a [subset of its functions](https://sites.google.com/site/bpoddocumentation/user-guide/function-reference), there is little indication of how they work together or how someone might be able to modify them.
- **Reproducibility** - As of [November 2020](https://github.com/pyControl/code/blob/cc6e7ab67c18388dea85b3ac48ac66a65ffa12f8/ChangeLog.txt#L74), pyControl has [versioned task files](https://pycontrol.readthedocs.io/en/latest/user-guide/pycontrol-data/#versioned-task-files) that append a hash to each version of a task and save it along with any produced data, tying the data to exactly the code that produced it. PyControl’s most recent releases have explicit [version numbers](https://github.com/pyControl/code/releases), but these don’t appear to be saved along with the data. Bpod stores neither code nor task versions in its data. Neither system saves experimental parameter changes by default—and the GUIs of both allow parameters to be changed at will—and so critical data could be lost and experiments made unreproducible unless the user writes custom code to save them. Bpod has an undocumented [plugin system](https://github.com/sanworks/Bpod_Gen2/commit/10ad997555086afb93dfc1080091acaa58d740f9), but neither system has a formal system for sharing plugins or task code, requiring work to be duplicated across all users of the system.
- **Integration and Extension** - Integration with other systems that might handle some out-of-scope function is tricky in both of these example systems. All systems have some limitation, so care must be taken to provide points by which other systems might interact with them. One particularly potent example is the use of Bpod in the International Brain Laboratory’s standardized experimental rig[27], which relies on a single-purpose [93 page PDF](https://figshare.com/articles/preprint/A_standardized_and_reproducible_method_to_measure_decision-making_in_mice_Appendix_3_IBL_protocol_for_setting_up_the_behavioral_training_rig/11634732) to describe how to use the [iblrig](https://github.com/int-brain-lab/iblrig) library, which consists of a large amount of single-purpose code for stitching together pybpod with [bonsai](https://github.com/int-brain-lab/iblrig/blob/18569278fc2d8cd3266adb2a5f660a43f8f2582e/iblrig/bonsai.py) for controlling video acquisition. Even if a system takes a large amount of additional work to integrate with another, hopefully the system allows it to be done in a way such that it can be reused and shared with others in the future so they can be spared the trouble. The relatively sparse [documentation](https://iblrig.readthedocs.io/en/latest/index.html) and the high proportion of ibl-specific code present in the repository make that seem unlikely.

**Figure 1.3:**
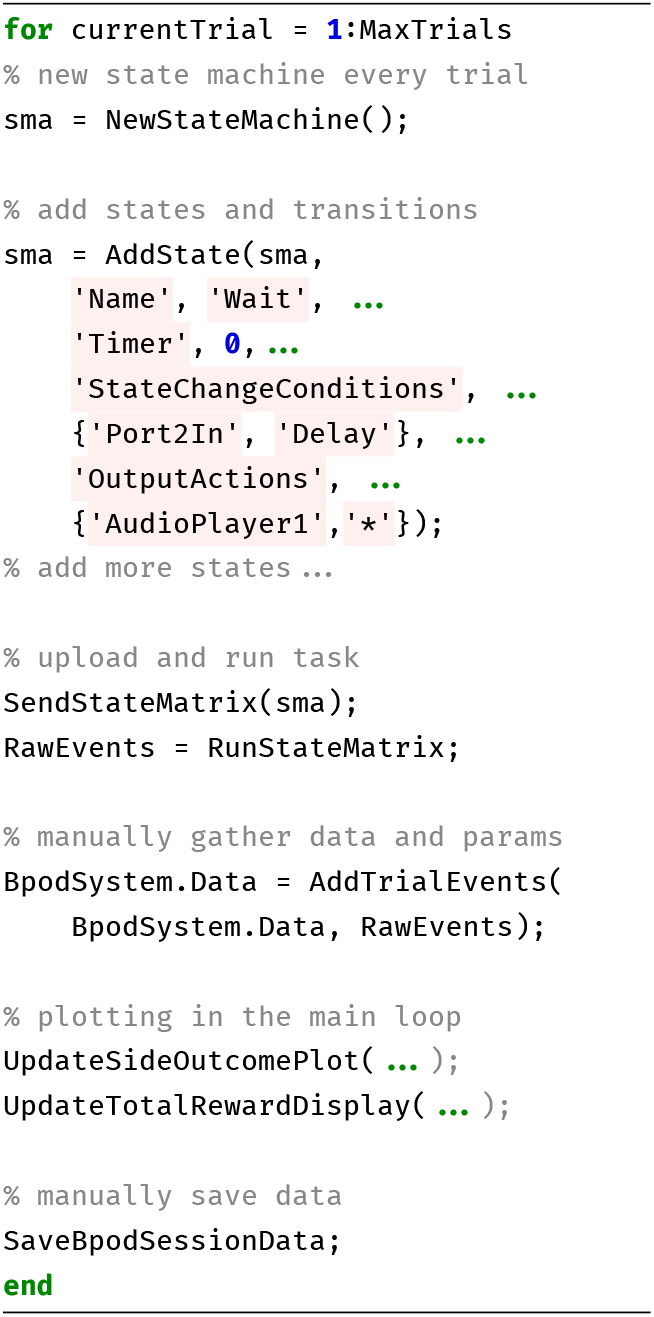
[Bpod’s general task structure](https://github.com/sanworks/Bpod_Gen2/blob/df6cd0c7d5df8247b02077b05fc263f79b86b096/Examples/Protocols/Sound/AnalogSound2AFC/AnalogSound2AFC.m).

Some of these limitations are cosmetic—fixable with additional code or hardware—but several of the most crucial are intrinsic to the design of these systems.

These systems, among others, have pioneered the development of modern behavioral hardware and software, and are to be commended for being open-source and highly functional. One need look no further for evidence of their usefulness than to their adoption by many labs worldwide. At the time that these systems were developed, a general-purpose single-board computer with performance like the Raspberry Pi 4 was not widely available. The above two systems are not unique in their limitations^7^, but are reflective of broader constraints of developing experimental tools: solving these problems is *hard*. We are only able to articulate the design principles that differentiate Autopilot by building on their work.

## 2 Design

Autopilot distributes experiments across a network of Raspberry Pis,^1^ a type of inexpensive single-board computer.

### Autopilot has three primary design principles

1. **Efficiency** - Autopilot should minimize computational overhead and maximize use of hardware resources.
2. **Flexibility** - Autopilot should be transparent in all its operations so that users can expand it to fit their existing or desired use-cases. Autopilot should provide clear points of modification and expansion to reduce local duplication of labor to compensate for its limitations.
3. **Reproducibility** - Autopilot should maximize system transparency and minimize the potential for the black-box of local reprogramming. Autopilot should maximize the information it stores about its operation as part of normal data collection.

### 2.1 Efficiency

Though it is a single board, the Raspberry Pi operates more like a computer than a microcontroller. It most commonly runs a custom Linux distribution, Raspbian, allowing Autopilot to use Python across the whole system. Using an interpreted language like Python running on Linux has inherent performance drawbacks compared to compiled languages running on embedded microprocessors. In practice these drawbacks are less profound than they appear on paper: Python’s overhead is negligible on modern processors^2^, jitter and performance can be improved by wrapping compiled code, etc. While we view the gain in accessibility and extensibility of a widely used high-level language like Python as outweighing potential performance gains from using a compiled language, Autopilot is nevertheless designed to maximize computational efficiency.

#### Concurrency

Most behavioral software is single-threaded (Figure 2.1), meaning the program will only perform a single operation at a time. If the program is busy or waiting for an input, other operations are blocked until it is finished.

**Figure 2.1:**
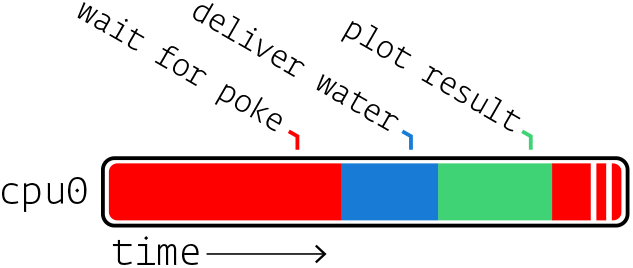
A single-threaded program executes all operations sequentially, using a single process and cpu core.

Autopilot distributes computation across multiple processes and threads to take advantage of the Raspberry Pi’s four CPU cores. Most operations in Autopilot are executed in **threads**. Specifically, Autopilot spawns separate threads to process messages and events, an architecture described more fully in section 3.8. Threading does not offer true concurrency^3^, but does allow Python to distribute computational time between operations so that, for example, waiting for an event does not block the rest of the program, and events are not missed because the program is busy (Figure 2.2).

**Figure 2.2:**
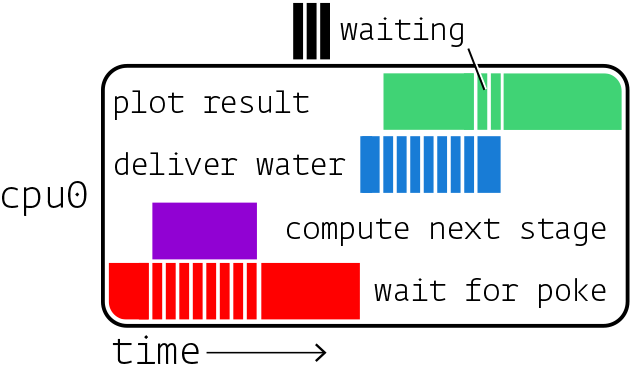
A multi-threaded program divides computation time of a single process and cpu core across multiple operations so that, for example, waiting for input doesn’t block other operations.

Critical operations that are computationally intensive or cannot be interrupted are given their own dedicated **processes**. Linux allows individual cores of a processor to be reserved for single processes, so individual Raspberry Pis are capable of running four truly parallel processing streams. For example, all Raspberry Pis in an Autopilot swarm create a messaging client to handle communication between devices which runs on its own processor core so no messages are missed. Similarly, if an experiment requires sound delivery, a realtime sound engine in a separate process (Figure 2.3) also runs on its own core.

**Figure 2.3:**
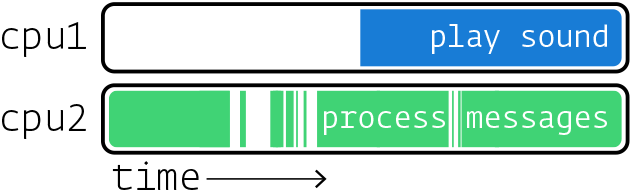
A multi-process program is truly concurrent, allowing multiple cpu cores to operate in parallel.

Since even moderately complex experiments can consume more resources than are available on a single processor, the topmost layer of concurrency in Autopilot is to use additional **computers**. Autopilot uses the Raspberry Pi as a low-cost hardware controller, but only its GPIO control system is unique to them: the rest of the code can be used on any type of computer, so computationally expensive or GPU-intensive operations can be offloaded to any number of high performance machines. Computers divide labor *autonomously* (see 2.2 and 3.7), so for example one computer running a task can send and receive messages from another running the GUI and plots, but does not *depend* on that input as it would in a system that couples a microcontroller with a managing computer. The ability to coordinate multiple, autonomous computers with heterogeneous responsibilities and capabilities in a shared task is Autopilot’s definitive design decision.

#### Leveraging Low-Level Libraries

Autopilot uses Python as a “glue” language, where it wraps and coordinates faster low-level compiled code[29]. Performance-critical components of Autopilot are thin wrappers around fast C libraries (Table 2.1). As Autopilot’s API matures, we intend to replace any performance-limiting Python code like its sound server and networking operations with compiled code exposed to python with tools like the C Foreign Function Interface ([CFFI](https://cffi.readthedocs.io/en/latest/index.html)).

**Table 2.1:** A few libraries Autopilot uses

|  |  |
| --- | --- |
| jack | realtime audio |
| pigpio | GPIO control |
| ZeroMQ | networking |
| Qt | GUI |

Since Autopilot coordinates its low-level components in parallel rather putting everything inside one “main loop,” Autopilot actually has *better* temporal resolution than single-threaded systems like Bpod or pyControl, despite the realtime nature of their dedicated processors (Table 2.2).

**Table 2.2:** Using pigpio as a dedicated I/O process gives autopilot greater measurement precision

|  | Precision |
| --- | --- |
| Autopilot (pigpio) | 5μs |
| Bpod | 100μs |
| pyControl | 1000μs |

#### Caching

Finite-state machines are only aware of the current state and the events that transition it to future states. They are thus incapable of exploiting the often predictable structure of behavioral tasks to precompute future states and precache stimuli. Further, to change task parameters between trials (eg. changing the rewarded side in a two-alternative forced-choice task), state machines need to be fully reconstructed and reuploaded to the device that runs them each time.

Autopilot precomputes and caches as much as possible. Rather than wait “inside” a state, Autopilot prepares each of the next possible events and saves them for immediate execution when the appropriate trigger is received. Static stimuli are prepared once at the beginning of a behavioral session and stored in memory. Before their presentation, they are buffered to minimize latency.

By providing full low-level documentation, we let researchers choose the balance between ease of use and performance themselves: it’s possible to just call a sound’s [play()](https://docs.auto-pi-lot.com/en/latest/stim/sound/base.html#autopilot.stim.sound.base.Jack_Sound.play) method, explicitly buffer it with its [buffer()](https://docs.auto-pi-lot.com/en/latest/stim/sound/base.html#autopilot.stim.sound.base.Jack_Sound.buffer) method, or generate samples on the fly with its play_continuous() method. Similarly, messages can be sent with a networking node’s [send()](https://docs.auto-pi-lot.com/en/latest/networking/node.html#autopilot.networking.node.Net_Node.send) method, or prepared beforehand by explicitly making a [Message](https://docs.auto-pi-lot.com/en/latest/networking/message.html#autopilot.networking.message.Message) and calling its [serialize()](https://docs.auto-pi-lot.com/en/latest/networking/message.html#autopilot.networking.message.Message.serialize) method.

Autopilot’s efficient design lets it access the best of both worlds—the speed and responsiveness of compiled code on dedicated microprocessors and the accessibility and flexibility of interpreted code.

### 2.2 Flexibility

#### Single-language

Behavior software that uses dedicated microprocessors must have some routine for compiling the high-level abstraction of the experiment into machine code. This gives those systems a theoretical advantage in processing speed, but the compiler becomes the bottleneck of complexity: only those things that can be compiled can be included in the experiment. This may in part contribute to the ubiquity of statemachine formalisms in behavior software.

Because Python is used throughout the system, extending Autopilot’s functionality is straightforward. Task design (see section 3.3) is effectively arbitrary—anything that can be expressed in Python is a valid task. This also allows Autopilot to easily be extended to make use of external libraries (eg. our integration with DeepLabCut-Live[30] and our planned integration with OpenEphys).

#### Modularity

Although Autopilot deeply integrates with the Raspberry Pi’s hardware, we have also worked to make its components modular. There is a tension between providing a full-featured behavioral system and the flexibility of its components — as additional features are added to a system, they can constrain the functionality of existing components that they rely on. To address this tension, we have continuously worked to decouple Autopilot into subcomponents with clear inheritance hierarchies and APIs that can used quasi-independently.

Modularity has 3 primary advantages:

1. **Modularity makes code more flexible** by reducing the constraints imposed by unstructured code interdependencies
2. **Modularity makes code more intelligible** by logically distributing tasks to discrete classes
3. **Modularity reduces effort-duplication** by allowing multiple, similar classes to be created with inheritance rather than copying and pasting.

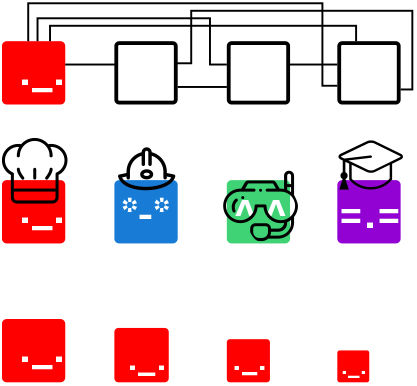

There is no such thing as “incompatible hardware” with Autopilot because the classes that control hardware are independent from the code that provides other core functionality. In systems without modular design, hardware implementation is spread across the codebase. For example to add a new type of hardware output to a Bpod system, one would need to write new firmware for it in C (eg. the [valve driver module](https://github.com/sanworks/Bpod_ValveDriver_Firmware)), [modify Bpod’s existing firmware](https://github.com/sanworks/Bpod_StateMachine_Firmware/blob/v22/Preconfigured/StateMachine-Bpod2_0/StateMachine-Bpod2_0.ino), hunt through the code to modify how [states are added](https://github.com/sanworks/Bpod_Gen2/blob/71f3a256b68926b65eae71e10fd747bd28e7ba7d/Functions/State%20Machine%20Assembler/AddState.m#L170) and state machines are assembled, add its controls explicitly [to the GUI](https://github.com/sanworks/Bpod_Gen2/blob/71f3a256b68926b65eae71e10fd747bd28e7ba7d/Functions/Override%20Panels/StateMachinePanel_2_0_0.m), and so on.

Tasks specify what type of hardware is needed to run them, but are agnostic about the way the hardware is implemented, making their descriptions more portable. Tasks that have the same structure but differ in hardware (eg. a freely moving two-alternativesystematic means of forced choice task in which a mouse visits several IR sensors, or a head-fixed two-alternative forced choice task in which a mouse runs on a wheel to indicate its choice) can be implemented by a trivial subclass that modifies the hardware description rather than completely rewriting the task.

#### Plugins & Code Transparency

We call Autopilot a software framework because in addition to providing classes and methods to run experiments out of the box, it also provides explicit structure that scaffolds any additional code that is needed by the user. Our goal is to clearly articulate in the documentation how modules should interact so that anyone can write code that works on any apparatus.

As groupware intended to be used differently by lab members with different responsibilities, Autopilot is designed for users with a range of programming expertise, from those who only want to interact with a GUI, to those who wish to fundamentally rewrite core operations for their particular experiment. As such, it is extensively documented: this paper provides a high-level introduction to its design and structure, its user guide describes how to use the program and provides examples, and its API-level documentation describes in granular detail how the program actually works^4^. Nothing is “off-limits” to the user—there isn’t any hidden, undocumented hardware code behind the curtain^5^. We want users to be able to understand how and why everything works the way it does so that Autopilot can be adapted and expanded to any use-case.

A broader goal of Autopilot is to build a library of flexible task prototypes that can be tweaked and adapted, hopefully reducing the number of times the wheel is reinvented. We have attempted to nudge users to write reusable tasks by designing Autopilot such that rather than writing tasks as local unstructured scripts, they use its plugin system that scaffolds development by extending any of its basic types. Plugins are registered using a form in the Autopilot Wiki which makes them [available to anyone](https://wiki.auto-pi-lot.com/index.php/Autopilot_Plugins) while also embedding them in a semantically annotated information system that allows giving explicit credit to contributors, programmatically linking to any derivative publications that use the plugin, and further documentation of any tasks, hardware, or other extensions included within the plugin. Inheriting from parent classes give plugins structure and a set of basic features^6^ while also being maximally permissive — anything can be overridden and modified.

#### Message Handling

Modular software needs a well-defined protocol to communicate between modules, and Autopilot’s is heavily influenced by the concurrency philosophy^7^ of ZeroMQ[31]. All communication between computers and modules happens with ZeroMQmessages, and handling those messages is the main way that Autopilot handles events. A key design principle is that Autopilot components should not “share state”—they can communicate, but they are not *dependent* on one another. While this may seem like a trivial detail, having networking and message-handling at its core has three advantages that make Autopilot a fundamental departure from previous behavioral software.

First, new software modules can be added to any system by simply dropping in a standalone networking object. There is no need to dramatically reorganize existing code to make room for new functionality. Instead new modules can receive, process, and send information by just connecting to another module in the swarm. For example, each plot opens a network connection to stream incoming task data independently from the stream that is saving the data.

Second, Autopilot can be made to interact with other software libraries that use ZeroMQ. For example, The OpenEphys GUI for electrophysiology [can send and receive ZMQ messages](https://open-ephys.atlassian.net/wiki/spaces/OEW/pages/23265310/Network+Events) to execute actions such as starting or stopping recordings. Interaction with other software is also useful in the case that some expensive computation needs to happen mid-task. For example, one could send frames captured from a video camera on a Raspberry Pi to a GPU computing cluster for tracking the position of the animal. Since ZeroMQ messages are just TCP packets it is also possible to communicate over the internet for remote control or to communicate with a data server.

Third, making every component network-capable allows tasks to be distributed over multiple Raspberry Pis. Chaining multiple Pis distributes the computational load, allowing, for example, one Raspberry Pi to record and process video while another runs a sound server and delivers rewards. Autopilot expands with the complexity of your task, simultaneously eliminating limitations on quantity of hardware peripherals while ensuring latency is minimal. More interestingly, distributing tasks allows the arbitrary construction of what we call “behavioral topologies,” which we describe in section 3.7.

### 2.3 Reproducibility

We take a broad view on reproducibility: including not only the ability to share data and recreate experiments, but also integrating into a broader ecosystem of tools that reduces labor duplication and encourages sharing and organizing technical knowledge. For us, reproducibility means building a set of tools that make every experiment and every technique available to anyone, anywhere.

#### Standardized task descriptions

The implementation and fine details of a behavioral experiment matter. Seemingly trivial details like milliseconds of delay between trial phases and microliters of reward volume can be the difference between a successful and unsuccessful task (Figure 2.4). *Reporting* those details can thus be the difference between a reproducible and unreproducible result. Researchers also often use “auxiliary” logic in tasks—such as methods for correcting response bias—that are never completely neutral for the interpretation of results. These too can be easily omitted due to brevity or memory in plain-English descriptions of a task, such as those found in Methods sections. Even if all details of an experiment were faithfully reported, the balkanization of behavioral software into systems peculiar to each lab (or even to individuals within a lab) makes actually performing a replication of a behavior result expensive and technically challenging. Widespread use of experimental tools that are not explicitly designed to preserve every detail of their operation presents a formidable barrier to rigorous and reproducible science[13].

**Figure 2.4:**
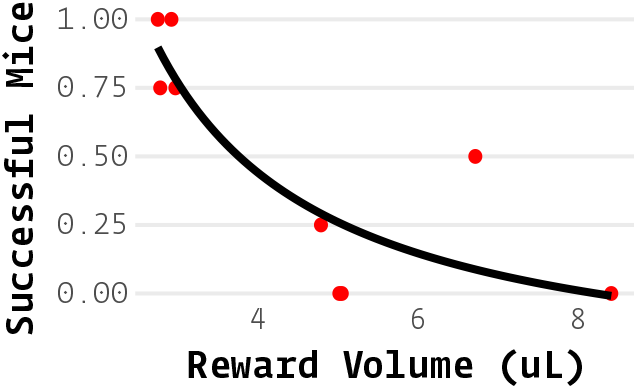
“Minor” details have major effects. Proportion of mice (each point, n=4) that were successful learning the first stage of the speech task described in [32] across 10 behavior boxes with variable reward sizes. A 2*μL* difference in reward size had a surprisingly large effect on success rate.

Autopilot splits experiments into a) the **code** that runs the experiment, which is intended to be standardized and shared across implementations, and b) the **parameters** (Figure 2.5) that define your particular experiment and system configuration. For example, two-alternative forced choice tasks have a shared structure regardless of the stimulus modality, but only your task plays pitch-shifted national anthems. This division of labor, combined with Autopilot’s structured plugin system, help avoid the ubiquitous problem of rig-specific code and hard-coded variables making experimental code only useful on the single rig it was designed for — enabling the possibility of a shared library of tasks as described in section 2.2

**Figure 2.5:**
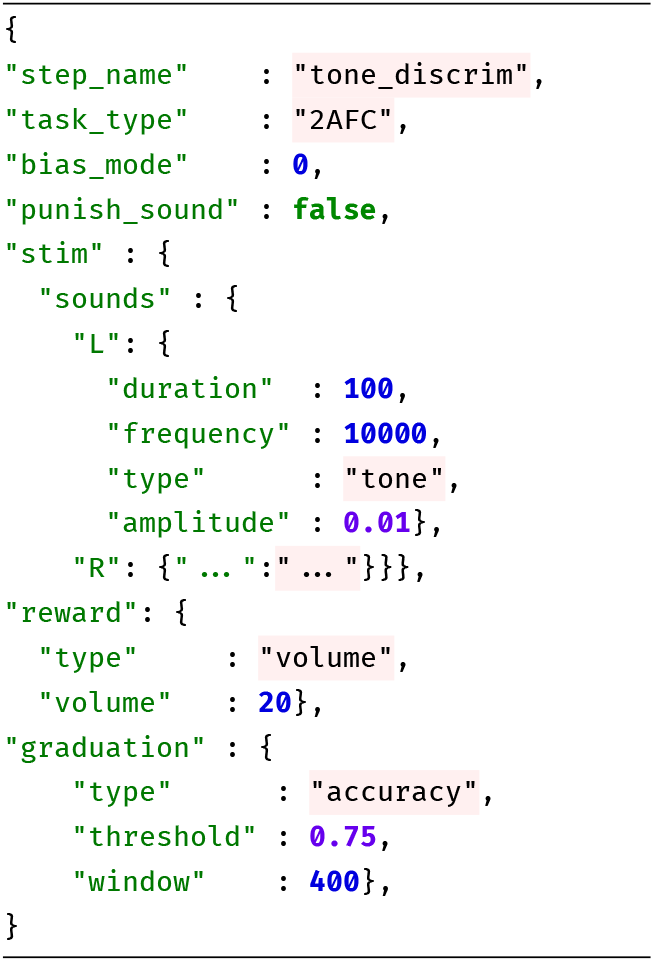
Task parameters are stored as portable JSON, formatting has been abbreviated for clarity.

The practice of reporting exactly the parameter description used by the software to run the experiment removes any chance for incompleteness in reporting. Because all task parameters are included in the produced data files, tasks are fully portable and can be reimplemented exactly by anyone that has comparable hardware to yours.

#### Self-Documenting Data

A major goal of the open science movement is to normalize publishing well documented and clearly formatted data alongside every paper. Typically, data are acquired and stored in formats that are lab-idiosyncratic or ad-hoc, which, over time, sprout entire software libraries needed just to clean and analyze it. Idiosyncratic data formats hinder collaboration within and between labs as the same cleaning and analysis operations gain multiple, mututally incompatible implementations, duplicating labor and multiplying opportunities for difficult to diagnose bugs. Over time these data formats and their associated analysis libraries can mutate and become incompatible with prior versions, rendering years of work inaccessible or uninterpretable. In one worst-case scenario, the cleaning process unearths some critically missing information about the experiment, requiring awkward caveats in the Methods section or months of extra work redoing it. In another, the missing information or bugs in analysis code are never discovered, polluting scientific literature with inaccuracies.

The best way to make data publishable is to avoid cleaning data altogether and *design good data hygiene practices into the data acquisition process*. Autopilot automatically stores all the information required to fully reconstruct an experiment, including any changes in task parameters or code version that happen throughout training as the task is refined.

Autopilot data is stored in HDF5 files, a hierarchical, high-performance file format. HDF5 files support metadata throughout the file hierarchy, allowing annotations to natively accompany data. Because HDF5 files can store nearly all commonly used data types, data from all collection modalities—trialwise behavioral data, continuous electrophysiological data, imaging data, etc.—can be stored together from the time of its acquisition. Data is always stored with the full conditions of its collection, and is ready to analyze and publish immediately (Figure 2.6). No Autopilot-specific scripts are needed to import data into your analysis tool of choice—anything that can read HDF5 files can read Autopilot data^8^.

**Figure 2.6:**
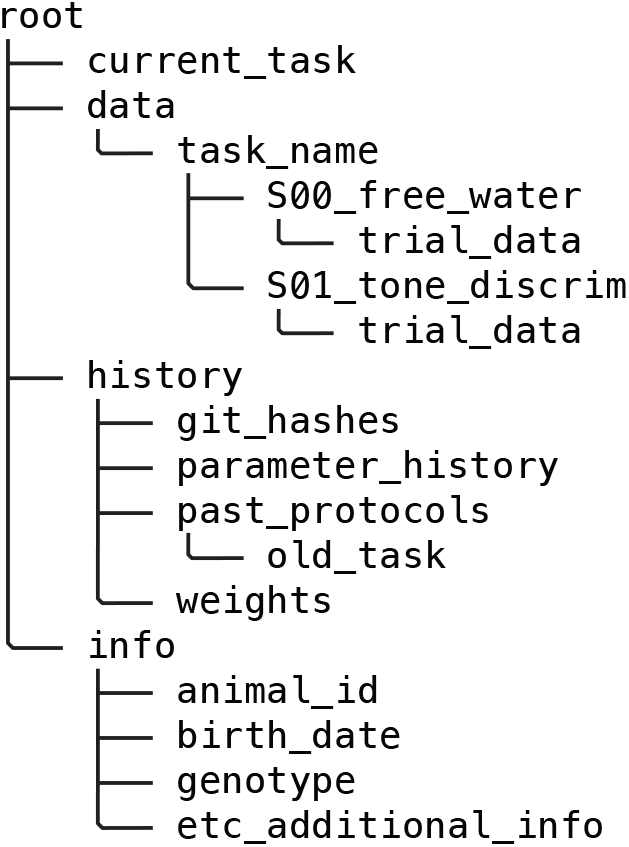
Example data structure. All information necessary to reconstruct an experiment is automatically stored in a human-readable HDF5 file.

As of v0.5.0, we have built a formal data modeling system into Autopilot, allowing for unified declaration of data for experimental subjects, task parameters, and resulting data with verifiable typing and human-readable annotations. These abstract data models can be used with multiple storage interfaces, paving the way for export to, for example, the Neurodata Without Borders standard[33], further enabling Autopilot data to be immediately incorporated into existing processing pipelines (see section 3.2).

#### Testing & Continuous Integration

Open-source scientific software does away with prior limitations to access and inspection imposed by proprietary tools. It also exposes the research process to bugs in software written by semi-amateurs that can yield errors in the resulting data, analysis, and interpretation[34, 35, 36, 37]. Autopilot tries to bring best practices in software development to experimental software, including a set of automated tests for continuous integration.

We are still formalizing our contribution process, and our tests are still far from achieving full coverage^9^, but we currently require tests and documentation for all new code added to the library. Writing good tests is hard, and we are in the process of building a set of hardware simulators and test fixtures to ease contribution.

Tests are effectively provable statements about how a program functions (Figure 2.7), which are particularly important for a library that aspires to be baseline lab infrastructure like Autopilot. Tests make it possible to use and contribute to the library with confidence: all tests are run on every commit, making it possible to determine if some new contribution breaks existing code without manually reading and testing every line. As we work to complete our test coverage, we hope to provide researchers with a tool that they can trust and elevates the verifiability of scientific results at large.

**Figure 2.7:**
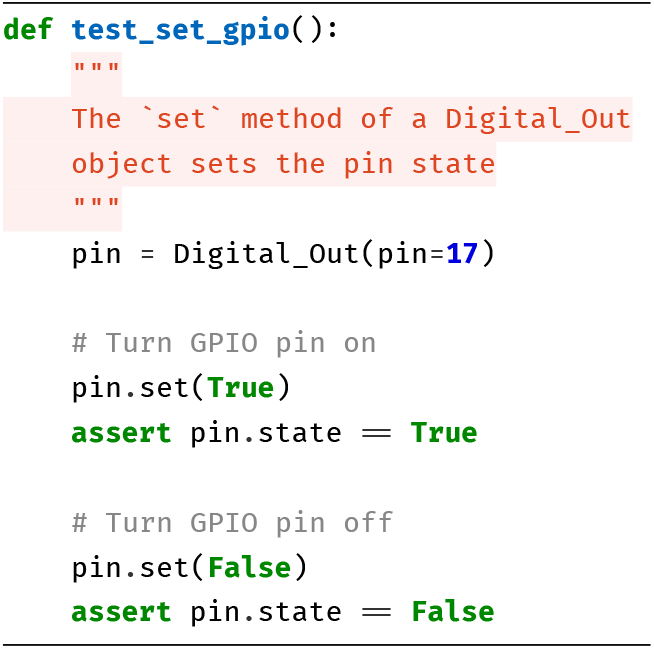
A test like `test_set_gpio` is a provable statement about the functionality of a program, in this case that “the `Digital_Out.set()` method sets the state of a GPIO pin.”

#### Expense

Autopilot is an order of magnitude less expensive than comparable behavioral systems (Table 2.3). We think the expense of a system is important for two reasons: scientific equity and statistical power.

**Table 2.3:**
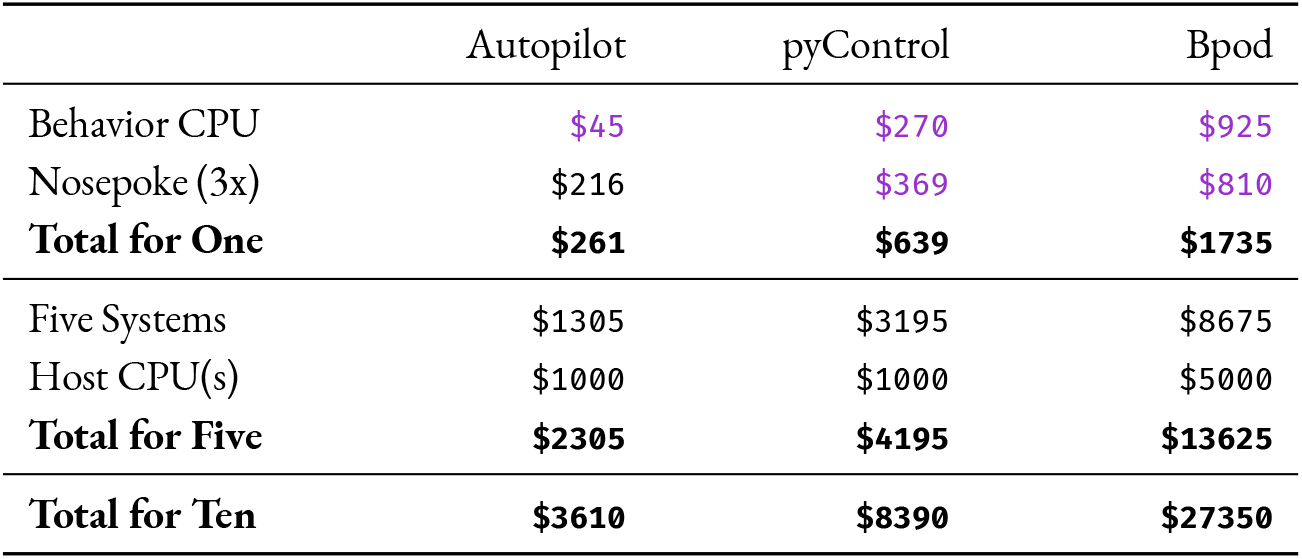
Cost for Basic 2AFC System. “Nosepoke” includes a solenoid valve, IR sensor, water tube, LED, housing, and any necessary driver PCBs. For PyControl and Autopilot, we included the cost of one Lee LHDA0531115H solenoid valve per nosepoke ($63.35). For PyControl, we estimated a typical USB hub with 5 ports to control 5 pyControl systems from one computer. We note that the Bpod and PyControl systems both include cost of assembly for the control CPUs and nosepokes, but also that Autopilot does not require assembly for its control CPU and its default nosepoke is a snap-together 3D printed part and PCB without surface mounted components that can be assembled by an amateur in roughly half an hour.

|  | Autopilot | pyControl | Bpod |
| --- | --- | --- | --- |
| Behavior CPU | \$45 | \$270 | \$925 |
| Nosepoke (3x) | \$216 | \$369 | \$810 |
| **Total for One** | **\$261** | **\$639** | **\$1735** |
| Five Systems | \$1305 | \$3195 | \$8675 |
| Host CPU(s) | \$1000 | \$1000 | \$5000 |
| **Total for Five** | **\$2305** | **\$4195** | **\$13625** |
| **Total for Ten** | **\$3610** | **\$8390** | **\$27350** |

The distribution of scientific funding is highly skewed, with a large proportion of research funding concentrated in relatively few labs[38]. Lower research costs benefit all scientists, but lower instrumentation costs directly increase the accessibility of state-of-the-art experiments to labs with less funding. Since well-funded labs also tend to be concentrated at a few (well-funded) institutions, lower research costs also broaden the base of scientists outside traditional research institutions that can stay at the cutting edge[39, 40, 41].

Neuroscience also stands to benefit from the lessons learned from the replication crisis in Psychology[43]. In neuroscience, underpowered experiments are the rule, rather than the exception[44]. Statistical power in neuroscience is arguably even worse than it appears, because large numbers of observations (eg. neural recordings) from a small number of animals are typically pooled, ignoring the nested structure of observations collected within individual animals. Increasing the number of cells recorded from a small number of animals dramatically increases the likelihood of Type I errors (Figure 2.8)—indeed, for values of within-animal correlation typical of neuroscientific data, high numbers of observations make Type I errors more likely than not[42]. For this reason, perhaps paradoxically, recent technical advances in multiphoton imaging and silicon-probe recordings will actually make statistical rigor in neuroscience *worse* if we don’t use analyses that account for the multilevel structure of the data and correspondingly record from the increased number of animals that they require.

**Figure 2.8:**
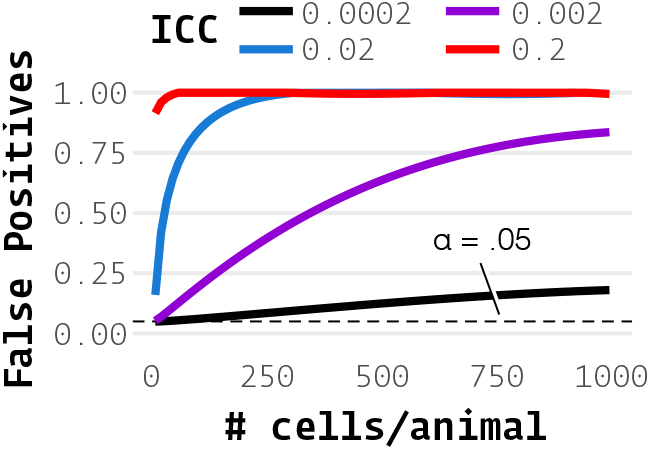
When comparing a value across groups, eg. a genetic knockout vs. wildtype, even a modest intra-animal (or, more generally, intra-cluster) correlation (ICC) causes the false positive rate to be far above the nominal *α* = 0.05. Shown are false positive rates for simulated data with various numbers of “cells” recorded for comparisons between two groups of 5 animals each with a real effect size of 0. We note that 741 simultaneously recorded cells were reported in [6] and a mean ICC of 0.19 across 18 neuroscientific datasets was reported in [42]

Although the expense of multi-photon imaging and high-density electrophysiology will always impose an experimental bottleneck, behavioral training time is often the greater determinant of study sample size. Typical behavioral experiments require daily training sessions often carried out over weeks and months, while far fewer imaging or electrophysiology sessions are carried out per animal. Training large cohorts of animals in parallel is thus the necessary basis of a well-powered imaging or electrophysiology experiment.

## 3 Program Structure

Autopilot consists of software and hardware modules configured to create a behavioral topology. Independent agents linked by flexible networking objects fill different roles within a topology, such as hosting the user interface, controlling hardware, transforming incoming and outgoing data, or delivering stimuli. This infrastructure is ultimately organized to perform a behavioral task.

### 3.1 Directory Structure

On setup, Autopilot creates a user directory that contains all local files that define its operation (Figure 3.2). The subdirectories include:

- **`calibration`** — Calibration for hardware objects like audio or solenoids that, for example, map opening durations to volumes of liquids dispensed
- **`data`** — Data for experimental subjects
- **`launch_autopilot.sh`** — Launch script that includes launching external processed like the jack audio daemon (will be removed and integrated into a more formal agent structure in future versions)
- **`logs`** — Every Autopilot object is capable of full debug logging, neatly formatted by object type and instance ID and grouped within module-level logging files. Logs are both written to disk, and output to stderr using the [rich logging handler](https://rich.readthedocs.io/en/latest/reference/logging.html#logging) for clean and readable inspection during program operation (Figure 3.3). Logs can be parsed back into python objects to make it straightforward to diagnose problems or recover data in the case of an error.
- **`pilot_db.json`** — A .json file that stores information about associated Pilots, including the contents of their prefs files, which hash/version of Autopilot they are running, and any Subjects that are associated with them.
- **`plugins`** — Plugins, which are any Python files that contain subclasses of Autopilot objects, that are automatically made available by Autopilot’s [registry](https://docs.auto-pi-lot.com/en/latest/utils/registry.html) system (eg. 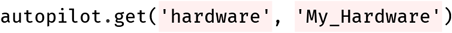 would retrieve a custom hardware object). Plugins can be documented and made available to other Autopilot users by registering them on the [wiki](https://wiki.auto-pi-lot.com/index.php/Autopilot_Plugins)
- **`prefs.json`** — Configuration options for this particular Autopilot instance, including configurations of local hardware objects, audio output, etc. In the future this will likely be broken into multiple files for different kinds of preferences^1^.
- **`protocols`** — Protocols, which consist of parameterizations of individual Tasks as well as criteria for graduating betwewen them. These are also stored in individual subject data files, and updated whenever the source protocol files change.
- **`sounds`** — Any sound files that are requested by the [File](https://docs.auto-pi-lot.com/en/latest/stim/sound/sounds.html#autopilot.stim.sound.sounds.File) sound class.

**Figure 3.1:**
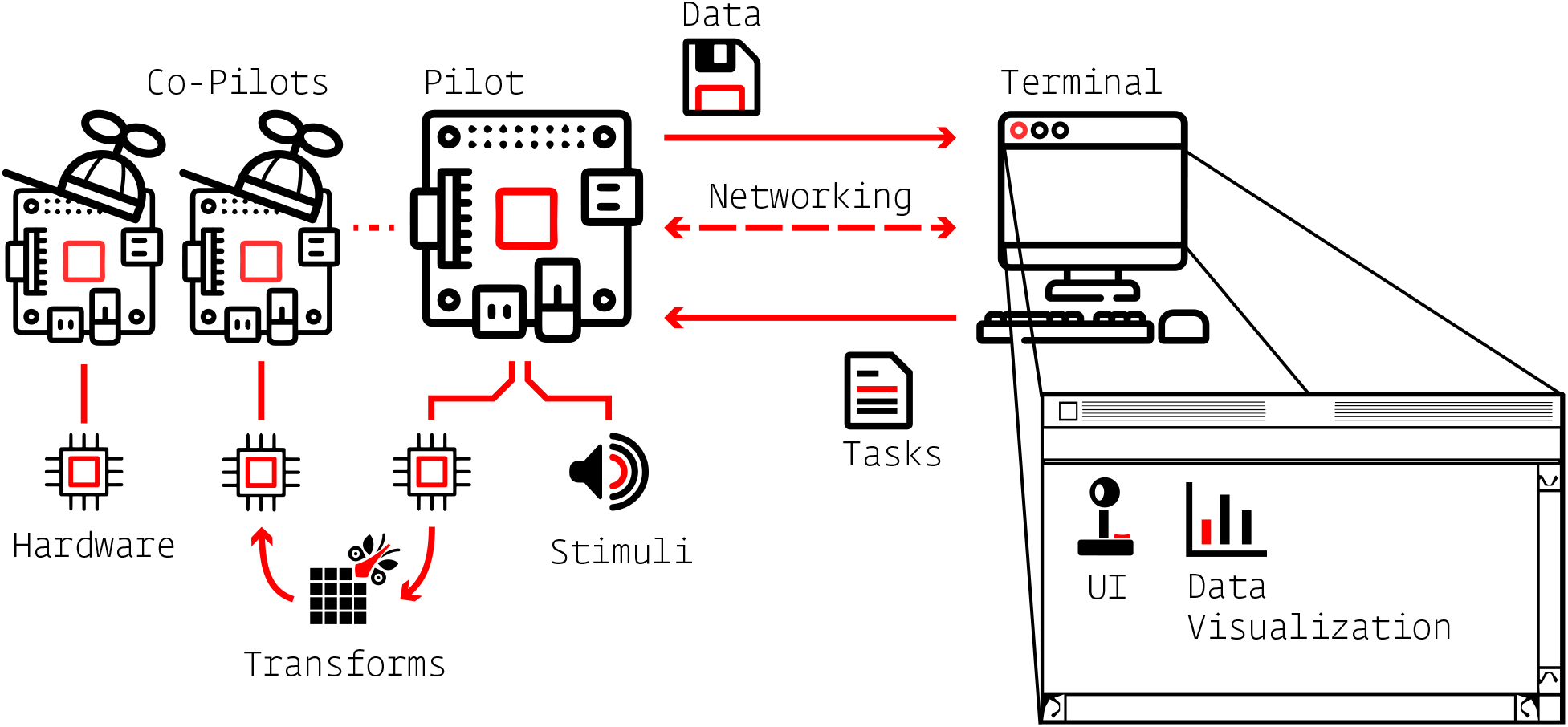
Overview of major Autopilot components

**Figure 3.2:**
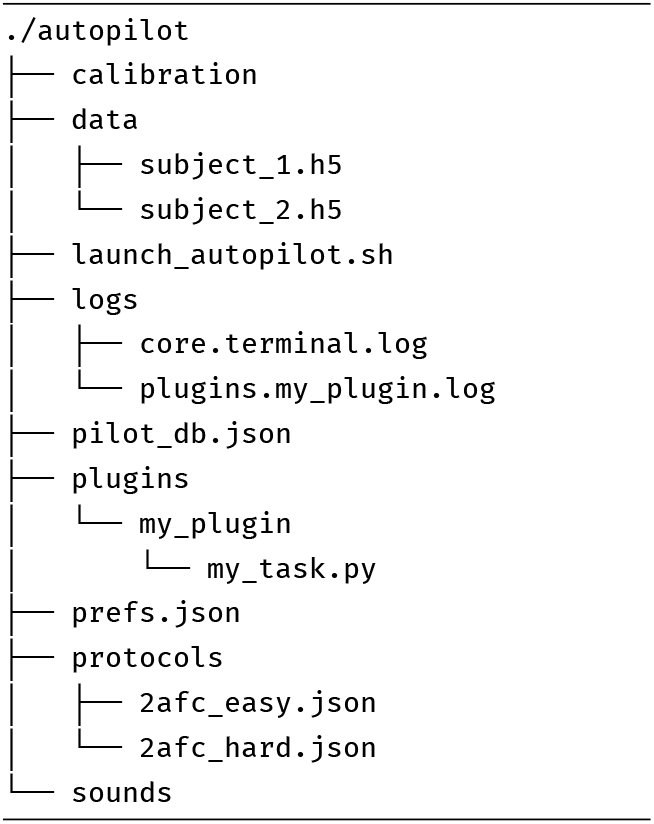
Example user directory structure, typically in `~/autopilot`.

**Figure 3.3:**
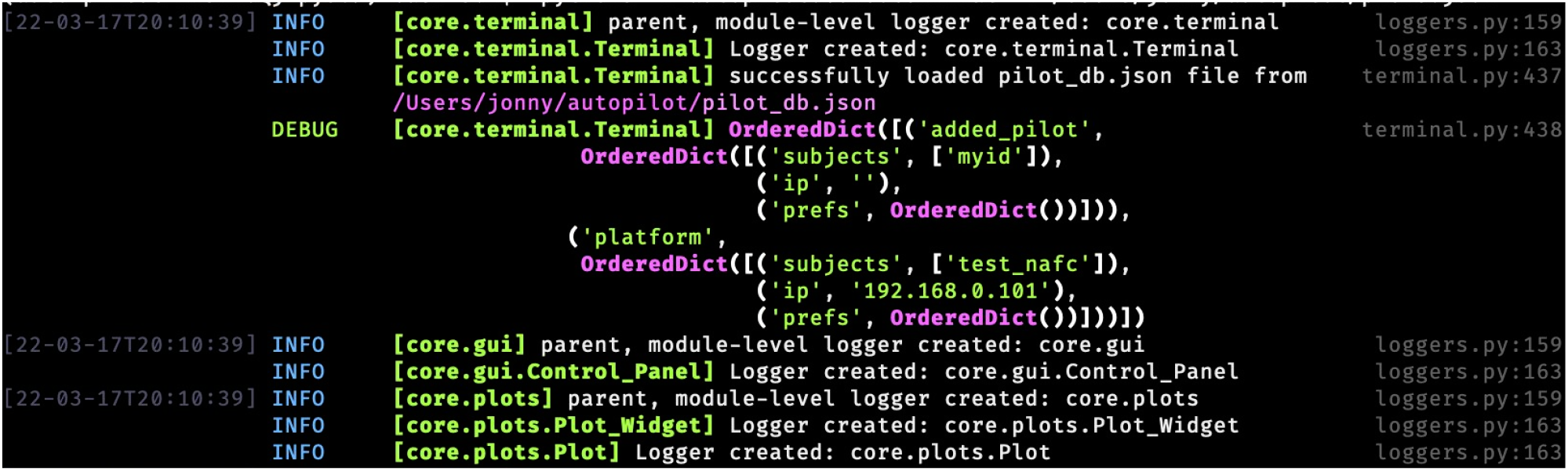
Logs printed to stderr are formatted and colorized by the [rich logging handler](https://rich.readthedocs.io/en/latest/reference/logging.html#logging). Logfiles are created by module, and log entries are identified by the individual objects instantiated from them. Logfiles are rotated and size-limited for configurable backups.

### 3.2 Data

As of v0.5.0, Autopilot uses [pydantic](https://pydantic-docs.helpmanual.io/) to create explicitly typed and schematized data models. Submodules include `data` abstract `modeling` tools that define base model types like `Tables`, `Groups`, and sets of `Attributes`. These base modeling classes are then built into a few core data models like subject `Biography` information, `Protocol` declaration, and the `Subject` data model itself that combines them. Modeling classes then have multiple `interfaces` that can be used to create equivalent objects in other formats, like pytables for hdf5 storage, pandas dataframes for analysis, or exported to Neurodata Without Borders.

For example, consider a simplified version of the Biography model:

**Listing 1:**
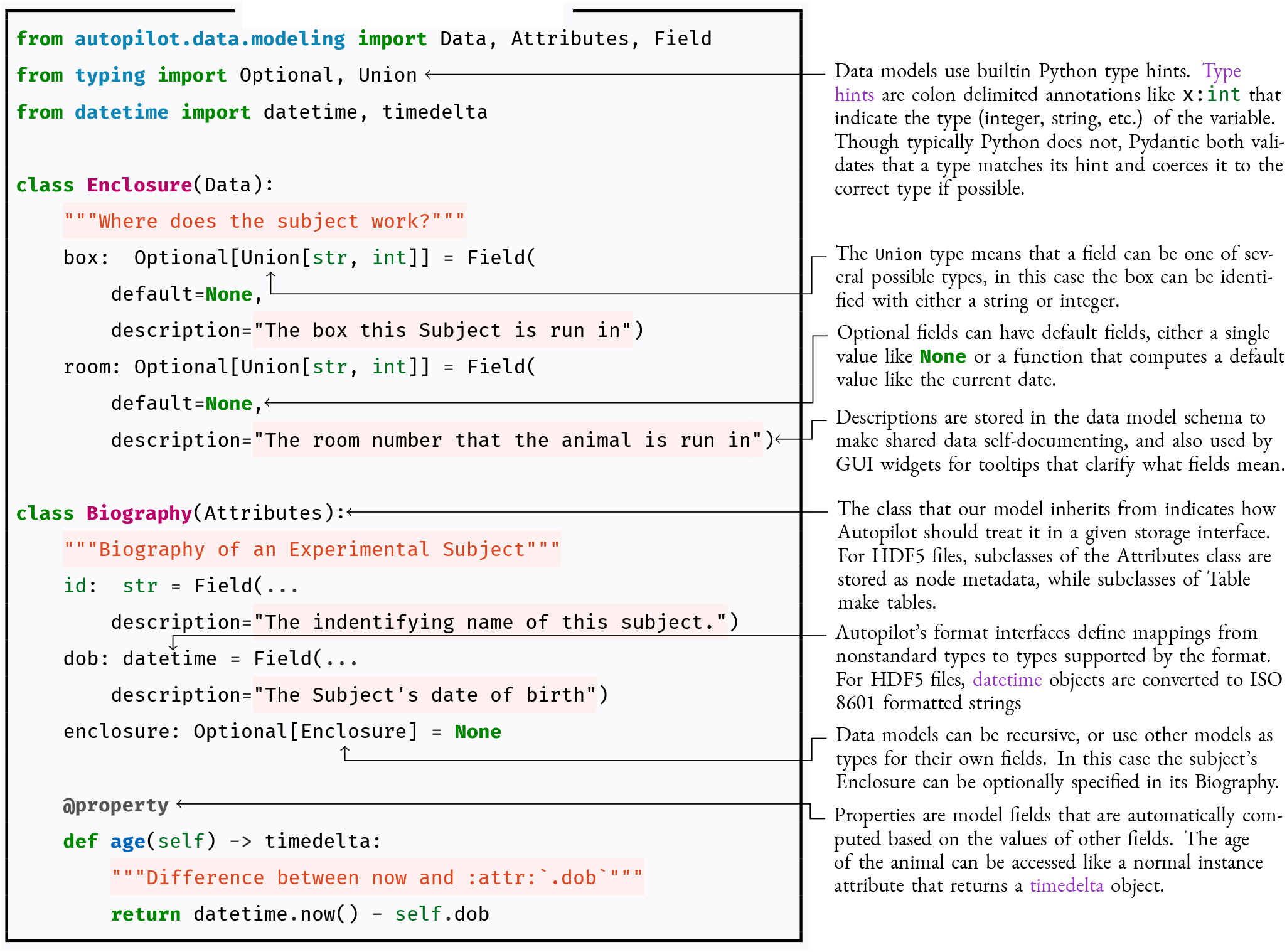
data - Biography.

A new subject could then be created with a biography like this, storing it in the HDF5 file and made accessible through the Subject interface:

**Listing 2:**
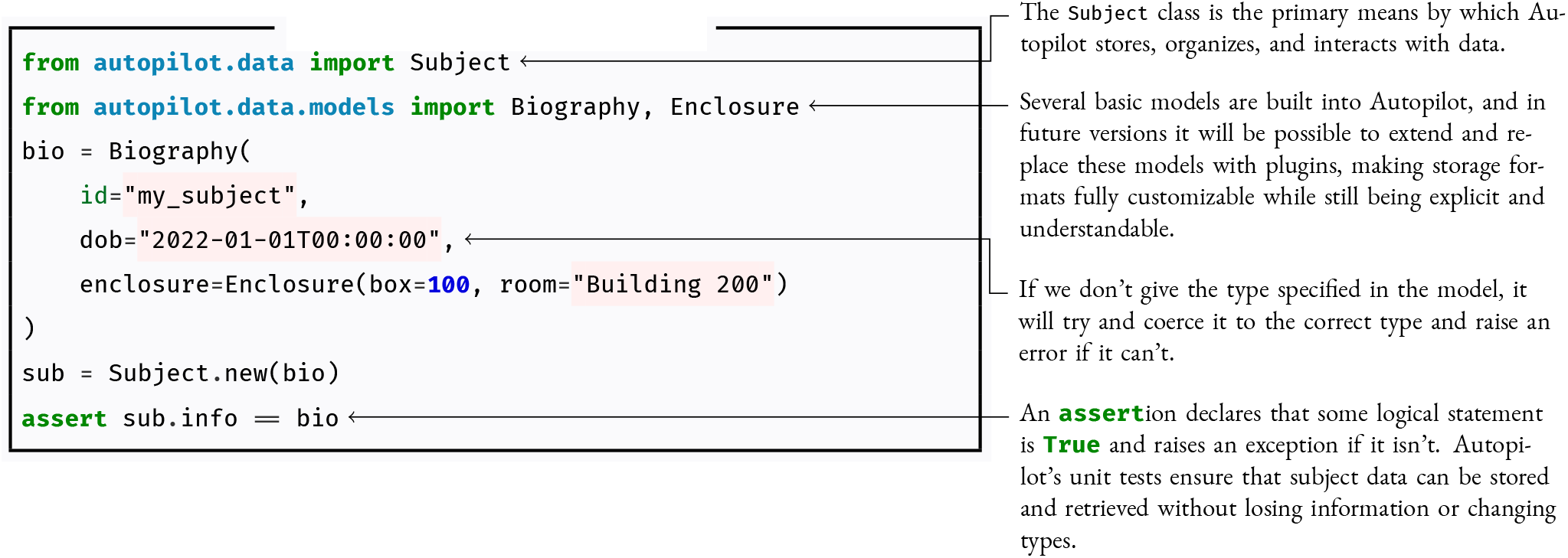
data - New Subject.

The models are declared using a combination of python type hints and `Field` objects that provide defaults and descriptions. Because these models can be recursive, as in the case of using the `Enclosure` model as a type within the `Biography` model, we can build expressive, flexible, but still strict representations of complex data.

Out of the box, pydantic models can create explicit and interoperable [schemas](https://pydantic-docs.helpmanual.io/usage/schema/) in [JSON Schema](https://json-schema.org/draft/2020-12/json-schema-core.html) and [OpenAPI](https://github.com/OAI/OpenAPI-Specification) formats, and Autopilot extends them with additional interfaces and representations. Autopilot can create a GUI form for filling in fields for models, for example, to create a new Subject or declare parameters for a task (Figure 3.4). `Attribute` models that consist of scalar key-value pairs can be reliably stored and retrieved from metadata attribute sets in HDF5 groups, but Autopilot knows that `Table` models should be created as HDF5 tables as they will have multiple values for each field. An additional `Trial_Data` class that inherits from `Table` can be exported to NWB trial data, and the `Subject.get_trial_data` method uses the model to load trial data and convert it to a correctly typed pandas[45] DataFrame.

**Figure 3.4:**
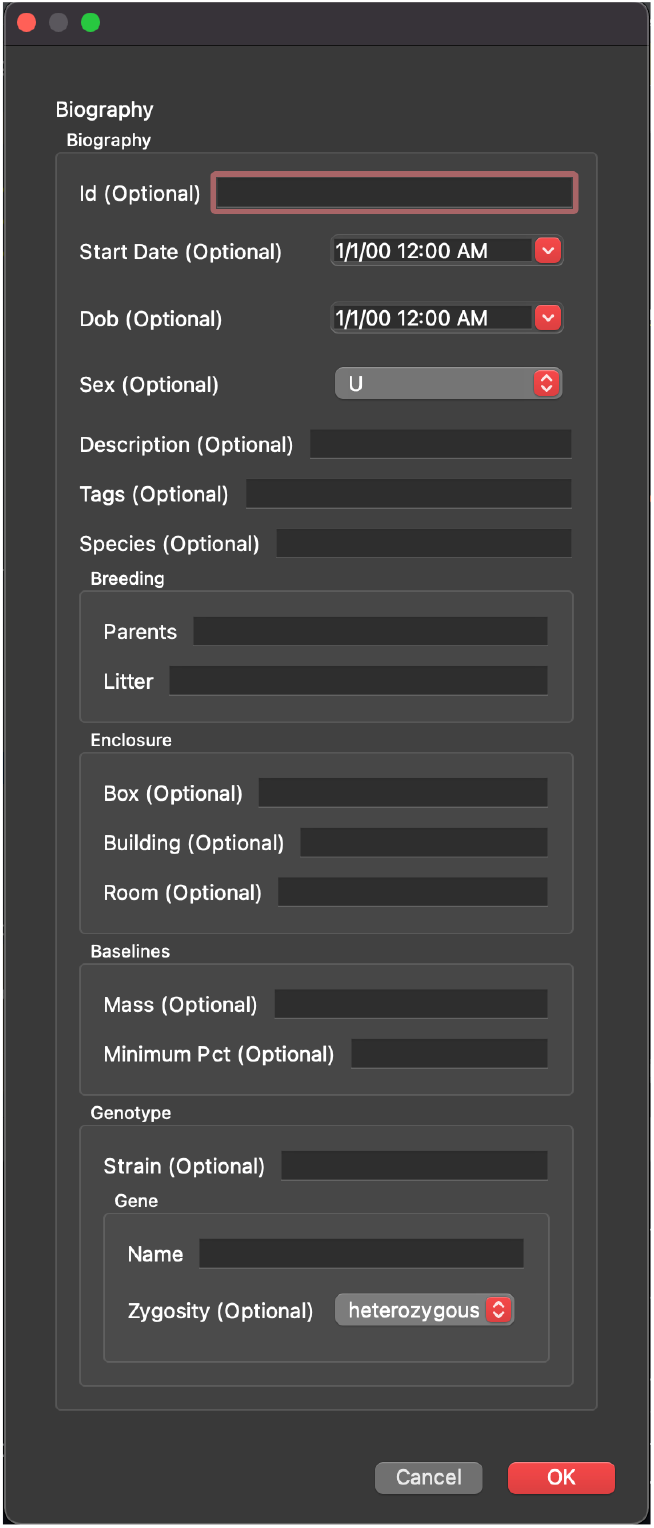
An Autopilot Data model can automatically generate a GUI form to fill in its properties, in this example to define a new experimental Subject’s biography.

Though the data modeling system is new in v0.5.0^2^, we have laid the groundwork for Autopilot’s plugin system to allow researchers to declare custom schema for all data produced by Autopilot, and to preserve both interoperability and reproducibility by combining them with datasets potentially produced by multiple incompatible tools (see Section 5.4).

### 3.3 Tasks

Behavioral experiments in Autopilot are centered around **tasks**. Tasks are Python classes that describe the parameters, coordinate the hardware, and perform the logic of the experiment. Tasks may consist of one or multiple **stages** like a stimulus presentation or response event, completion of which constitutes a **trial** (Figure 3.5). Stages are analogous to states in the finite state machine formalism.

**Figure 3.5:**
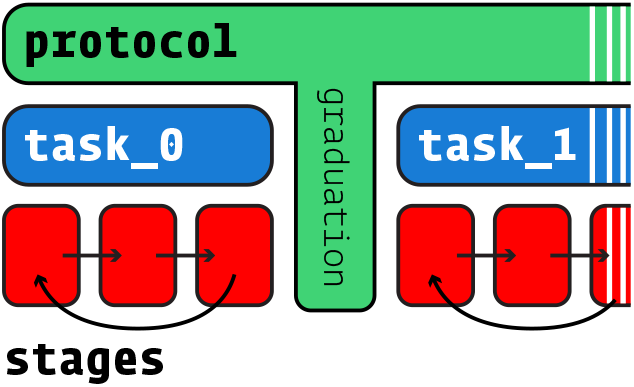
Protocols consist of one or multiple tasks, tasks consist of one or multiple stages. Completion of all of a task’s stages constitutes a trial, and meeting some graduation criterion like accuracy progresses a subject between tasks.

Multiple tasks are combined to make **protocols**, in which animals move between tasks according to “graduation” criteria like accuracy or number of trials. Training an animal to perform a task typically requires some period of shaping where they are familiarized to the apparatus and the structure of the task. For example, to teach animals about the availability of water from “nosepoke” sensors, we typically begin with a “free water” task that simply gives them water for poking their nose in them. Having a structured protocol system prevents shaping from relying on intuition or ad hoc criteria.

#### Task Components

The following is a basic two-alternative choice (2AFC) task—a sound is played and an animal is rewarded for poking its nose in a designated target nosepoke. While simple, it is included here in full to show how one can program a task, including an explicit data and plotting structure, in roughly 60 lines of generously spaced Python.

Every task begins by describing four elements:

1) the task’s parameters, 2) the data that will be collected, 3) how to plot the data, and 4) the hardware that is needed to run the task.

**Listing 3:**
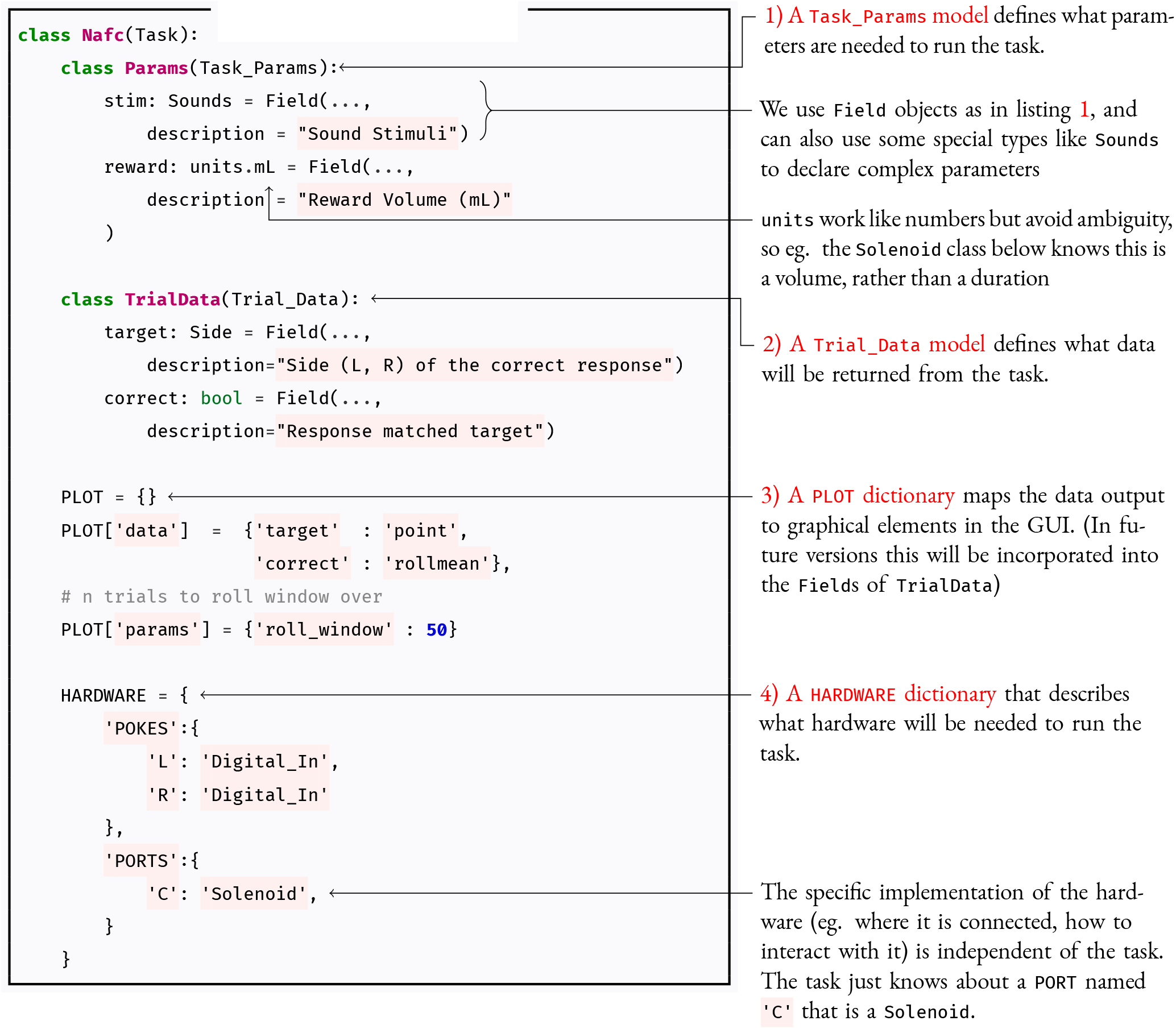
task - parameters.

Created tasks receive some common methods, like input/trigger handling and networking, from an inherited metaclass. Python inheritance can also be used to make small alterations to existing tasks^3^ rather than rewriting the whole thing. The GUI will use the `Params` model and the PLOT dictionary to generate forms for parameterizing the task within a protocol and display the data as it is collected. The `Subject` class will use the `TrialData` model to create HDF5 tables to store the data, and the `Task` metaclass will instantiate the described `HARDWARE` objects from their systemspecific configuration in the `prefs.json` file so they are available in the rest of the class like 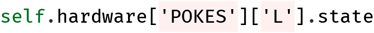

#### Stage Methods

The logic of tasks is described in one or a series of methods (stages). The order of stages can be cyclical, as in this example, or can have arbitrary logic governing the transition between stages.

**Listing 4:**
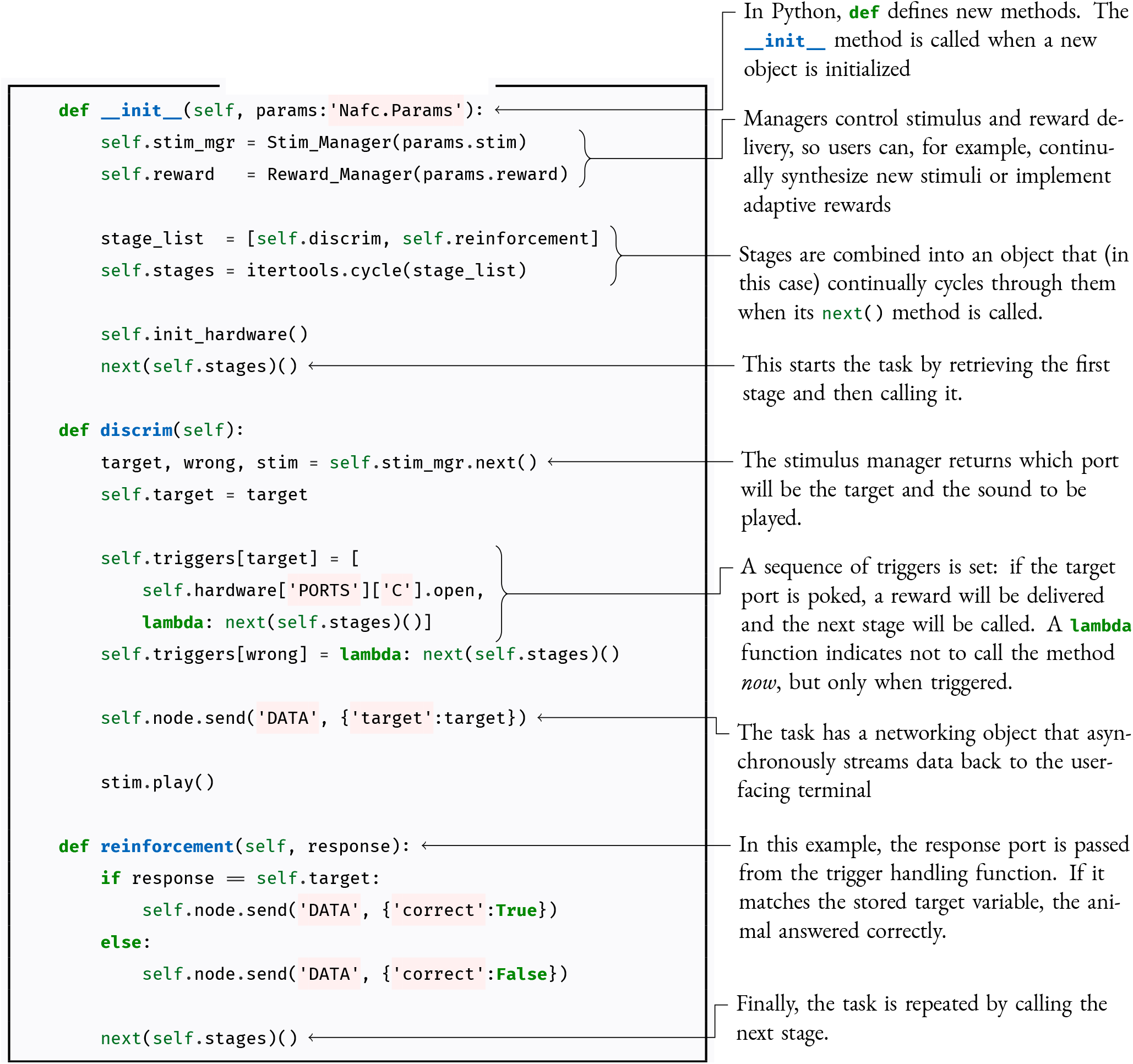
task - methods.

Autopilot is not prescriptive about how tasks are written. The same task could have two separate methods for correct and incorrect answers rather than a single reinforcement method, or only a single stage that blocks the program while it waits for a response.

Publishing data from this task requires no additional effort: a hash that uniquely identifies the code version (as well as any local changes) is automatically stored at the time of collection, as is a JSON-serialized version of the parameter model (Figure 3.6). If this task was incorporated into the central task library, anyone using Autopilot would be able to exactly replicate the experiment from the published data.

**Figure 3.6:**
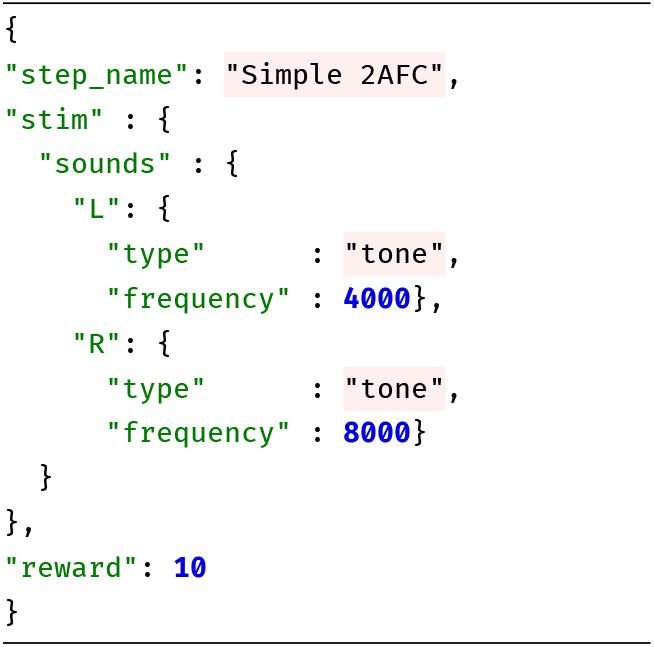
Simplified example of parameters for the above task

#### The limitations of finite state machines

The 2AFC task described above could be easily implemented in a finite-state machine. However, the difficulty of programming a finite-state machine is subject to combinatoric explosion with more complex tasks. Specifically, finite-state machines can’t handle any task that requires any notion of “state history.”

As an example, consider a maze-based task. In this task, the animal has to learn a particular route through a maze—it is not enough to reach the endpoint, but the animal has to follow a specific path to reach it (Figure 3.7). The arena is equipped with an actimeter that detects when the animal enters each area.

**Figure 3.7:**
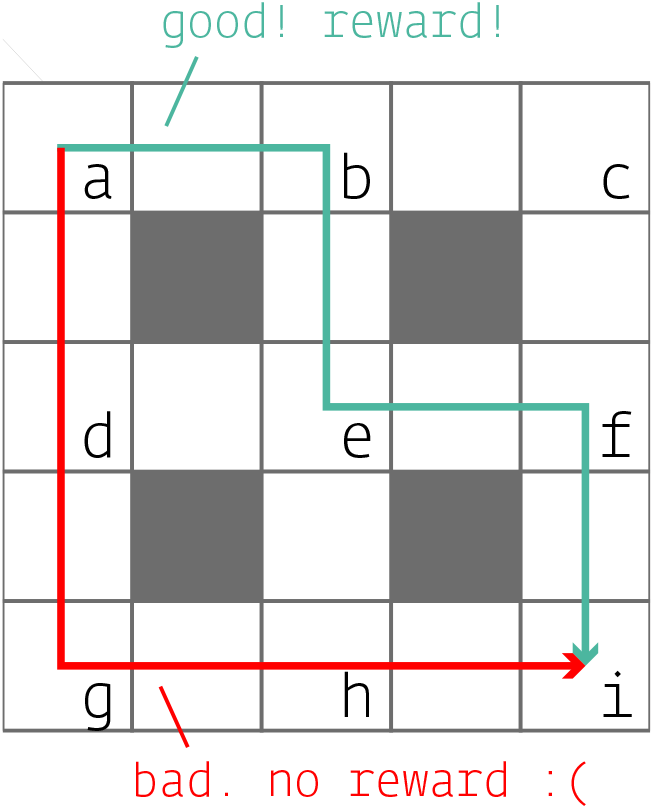
The subject must reach point i but only via the correct (green) path.

In Autopilot, we would define a hardware object that logs positions from the actimeter with a `store_position()` method. If the animal has entered the target position (“i” in this example), a `task_trigger()` that advances the task stage is called. The following code is incomplete, but illustrates the principle.

**Listing 5:**
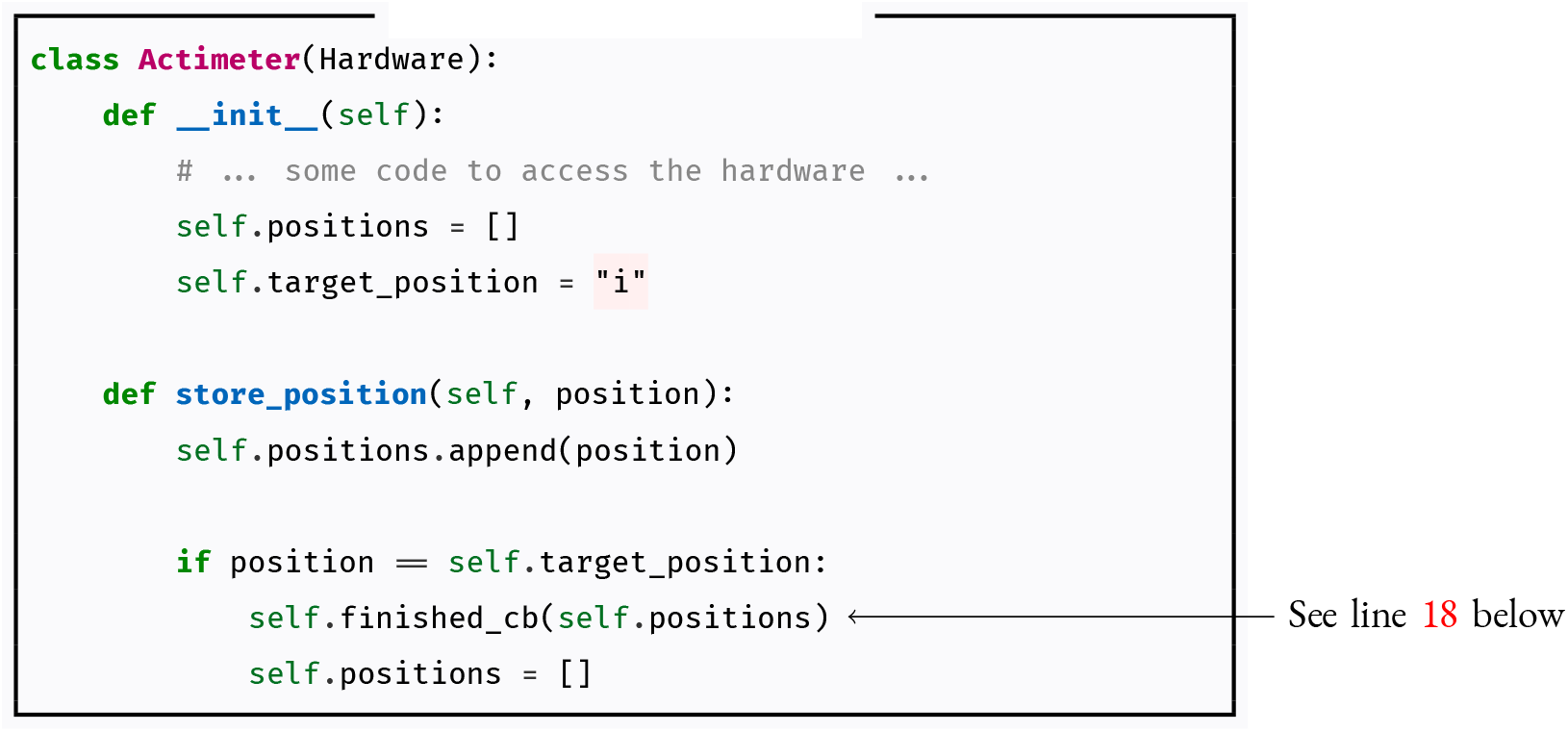
maze - hardware.

The task follows, with parameters and network methods for sending data omitted for clarity.

**Listing 6:**
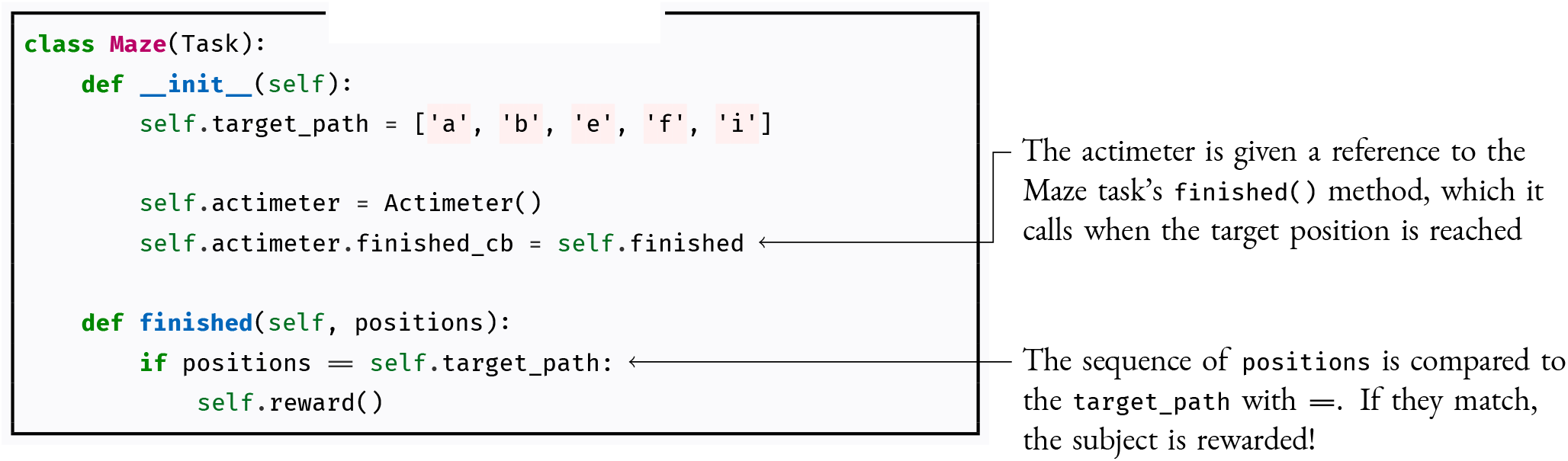
maze - task.

How would such a task be programmed in a finite-state machine formalism? Since the path matters, each “state” needs to consist of the current position and all the positions before it. But, since the animal can double back and have arbitrarily many state transitions before reaching the target corner, this task is impossible to represent with a finite-state machine, as a full representation would necessitate infinitely many states (this is one example of the *pumping lemma*, see [46]).

Even if we dramatically simplify the task by 1) assuming the animal never turns back and visits a space twice, and 2) only considering paths that are less than or equal to the length of the correct path, the finite state machine would be as complex as figure 3.8.

**Figure 3.8:**
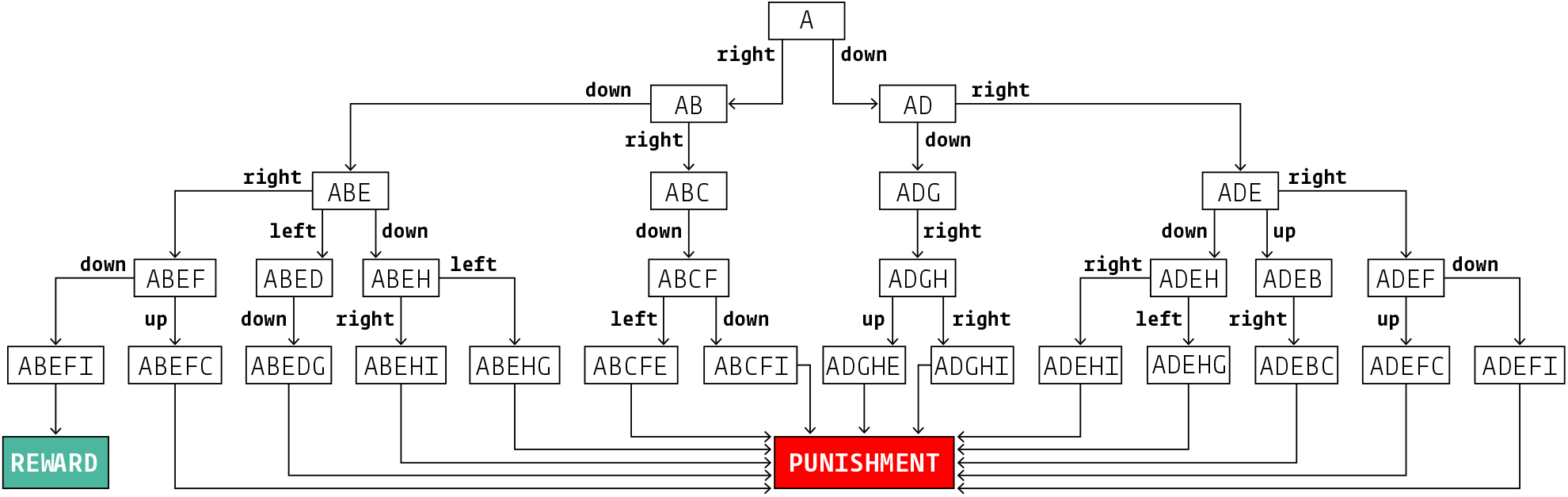
State transition tree for a simplified maze task.

**Figure 3.9:**
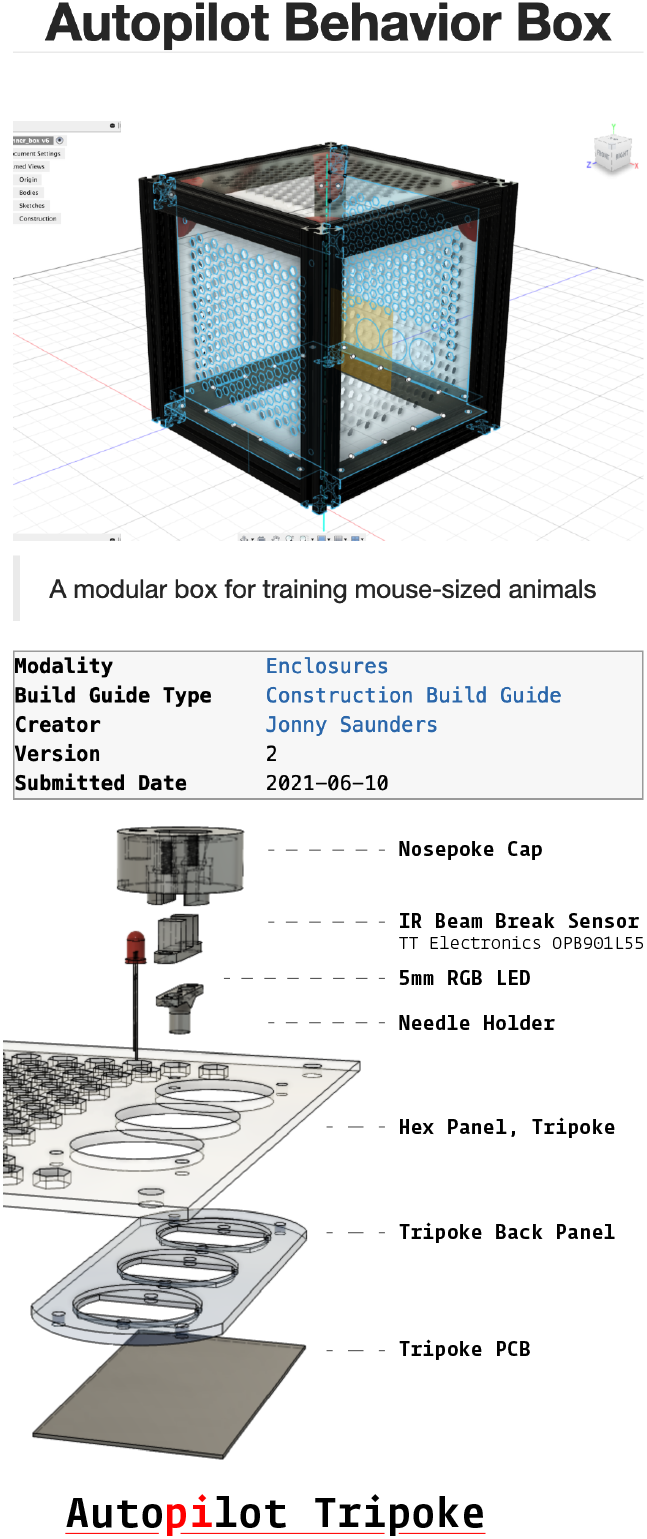
Two examples of parts with assembly guides available on the autopilot wiki: A modular [behavior box](https://wiki.auto-pi-lot.com/index.php/Autopilot_Behavior_Box) with magnetic snap-in panels (top), and a [three-nosepoke panel](https://wiki.auto-pi-lot.com/index.php/Autopilot_Tripoke) (bottom).

While finite-state machines are relatively easy to implement and work well for simple tasks, they quickly become an impediment to even moderately complex tasks. Even for 2AFC tasks, many desirable features are difficult to implement with a finite state machine, such as: (1) graduation to a more difficult task depending on performance history, (2) adjusting reward volume based on learning rate, (3) selecting or synthesizing upcoming stimuli based on patterns of errors[47], etc.

Some of these problems are avoidable by using extended versions of finite state machines that allow for extra-state logic, but require additional complexity in the code running the state machines to accomodate, and with enough exceptions the clean systematicity that is the primary benefit of finite state machines is lost. Autopilot attempts to avoid these problems by providing *tools* to program tasks and describe them without *requiring a specified format*, balancing the increased complexity by scaffolding the broader ecosystem of the experiment like its output data, hardware control, etc. When possible, we have tried to avoid forcing people to change the way they think about their work to fit our “little universe”^4^ and instead try to provide a set of tools that let researchers decide how they want to use them.

### 3.4 Hardware

The Raspberry Pi can interface with nearly all common hardware, and has an [extensive collection](https://www.raspberrypi.org/help/) of [guides](https://elinux.org/RPi_Guides), [tutorials](https://elinux.org/RPi_Tutorials), and an active [forum](https://www.raspberrypi.org/forums/) to support users implementing new hardware. There is also an enormous amount of existing hardware for the Raspberry Pi, including [sound cards](https://www.hifiberry.com/), [motor controllers](https://www.adafruit.com/product/2348), [sensor arrays](https://www.digikey.com/product-detail/en/raspberry-pi/SENSE-HAT/1690-1013-ND/6196429), [ADC/DACs](https://www.seeedstudio.com/Raspberry-Pi-High-Precision-AD-DA-Board-p-2765.html), and [touchscreen displays](https://www.digikey.com/product-detail/en/pimoroni-ltd/PIM369/1778-1221-ND/9521981), largely eliminating the need for a separate ecosystem of purpose-built hardware (Table 3.1).

**Table 3.1:** Cost of common peripherals. The native hardware of the Raspberry Pi, low-level hardware control of Autopilot, and availability of inexpensive off-the-shelf components compatible with the raspi make most custom-built peripherals unnecessary.

| Device | Raspi | Bpod |
| --- | --- | --- |
| HiFiBerry DAC2 Pro | \$45 | \$445 |
| ADC | \$30 | \$495 |
| I2C | \$0 | \$225 |
| Ethernet | \$0 | \$285 |
| Rotary Encoder | \$0 | \$145 |

**Table 3.2:**
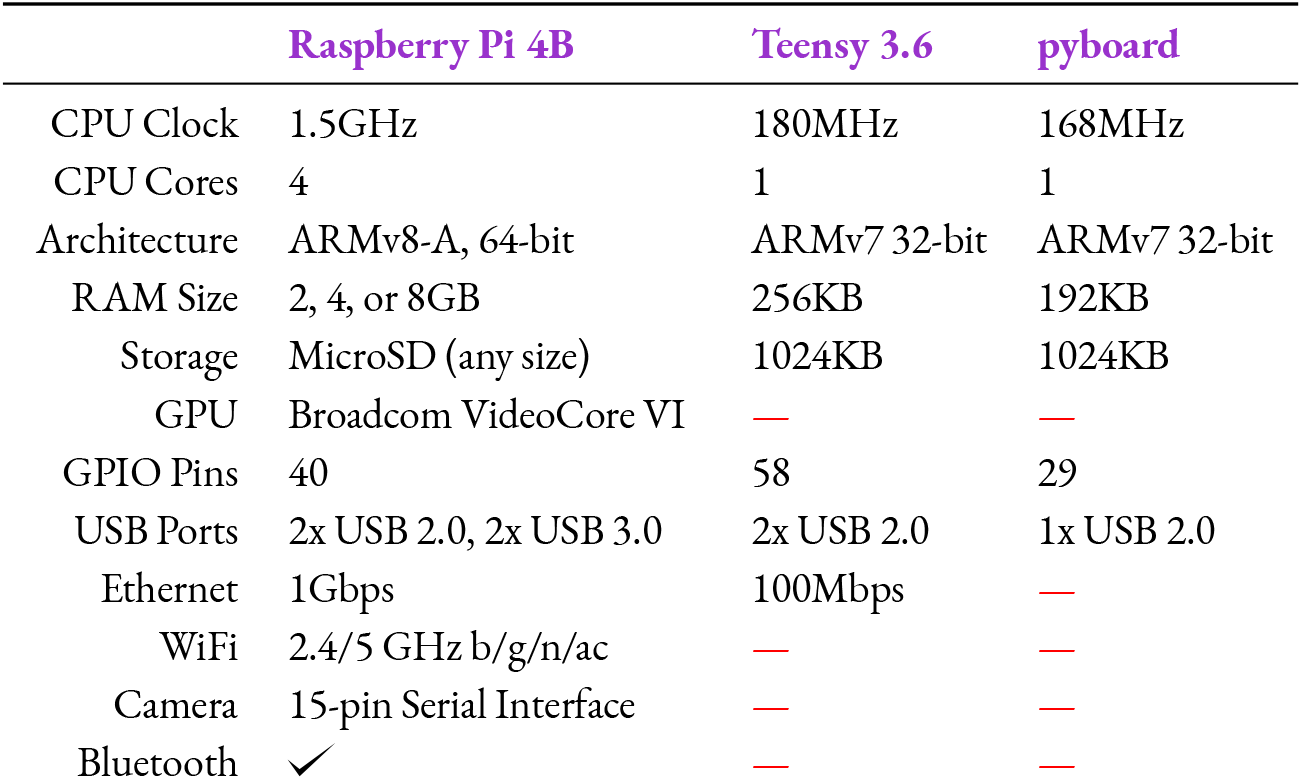
Specifications of reviewed behavior hardware. BPod’s state machine uses the Teensy 3.6 microcontroller, and PyControl uses the Micropython Pyboard.

Autopilot controls hardware with an extensible inheritance hierarchy of Python classes intended to be built into a library of hardware controllers analogously to tasks. Autopilot uses [pigpio](http://abyz.me.uk/rpi/pigpio/) to interact with its GPIO pins, giving Autopilot 5*μs* measurement precision and enabling protocols that require high precision (such as Serial, PWM, and I2C) for nearly all of the pins. Currently, Autopilot also has a family of objects to control cameras (both the [Raspberry Pi Camera](https://docs.auto-pi-lot.com/en/latest/hardware/cameras.html#autopilot.hardware.cameras.PiCamera) and [high-speed GENICAM-compliant](https://docs.auto-pi-lot.com/en/latest/hardware/cameras.html#autopilot.hardware.cameras.Camera_Spinnaker) cameras), i2c-based [motion](https://docs.auto-pi-lot.com/en/latest/hardware/i2c.html#autopilot.hardware.i2c.I2C_9DOF) and [heat](https://docs.auto-pi-lot.com/en/latest/hardware/i2c.html#autopilot.hardware.i2c.MLX90640) sensors, and [USB mice](https://docs.auto-pi-lot.com/en/latest/hardware/usb.html#autopilot.hardware.usb.Wheel). In the future we intend to improve performance further by replacing time-critical hardware operations with low-level interfaces written in Rust.

To organize and make available the vast amount of contextual knowledge needed to build and use experimental hardware, we have made a densely linked and publicly editable [semantic wiki](https://wiki.auto-pi-lot.com/index.php/Autopilot_Wiki). The Autopilot wiki contains, among others, reference information for [off-the-shelf parts](https://wiki.auto-pi-lot.com/index.php/Parts), schematics for [2D](https://wiki.auto-pi-lot.com/index.php/2D_CAD) and [3D](https://wiki.auto-pi-lot.com/index.php/3D_CAD)-printable components, and [guides](https://wiki.auto-pi-lot.com/index.php/Guides) for building experimental apparatuses and custom parts. The wiki combines unrestrictive freeform editing with structured, computer-readable [semantic properties](https://www.semantic-mediawiki.org/wiki/Semantic_MediaWiki), and we have defined a collection of [schemas](https://wiki.auto-pi-lot.com/index.php/Special:CategoryTree?target=Hardware&mode=categories&namespaces=&title=Special%3ACategoryTree) for commonly documented items coupled with [submission forms](https://wiki.auto-pi-lot.com/index.php/Special:Forms) for ease of use. For example, the wiki page for the [Lee Company solenoid](https://wiki.auto-pi-lot.com/index.php/Lee_LHDA0531115H) we use has fields from a generic [Part](https://wiki.auto-pi-lot.com/index.php/Category:Part) schema like a datasheet, price, and voltage, but also that it’s a 3-way, normally-closed [solenoid](https://wiki.auto-pi-lot.com/index.php/Category:Solenoid).

The wiki’s blend of structure and freedom breaks apart typically monolithic hardware documentation into a collaborative, multimodal technical knowledge graph. Autopilot can access the wiki through its [API](https://docs.auto-pi-lot.com/en/latest/guide/plugins.html#the-wiki-api), and we intend to tighten their integration over time, including automatic configurations for common parts, usage and longevity benchmarks, detecting mutually incompatible parts, and automatically resolving any additional plugins or dependencies needed to use a part.

### 3.5 Transforms

In v0.3.0, we introduced the `transform` module, a collection of tools for transforming data. The raw data off a sensor is often not in itself useful for performing an experiment: we want to compare it to some threshold, extract positions of objects in a video feed, and so on. Transforms are like building blocks, each performing some simple operation with a standard object structure, and then composed into a pipeline (Figure 3.10). Pipelines are portable, and can be created on the fly from a JSON representation of their arguments, so it’s easy to offload expensive operations to a more capable machine for distributed realtime experimental control (See [30]).

**Figure 3.10:**
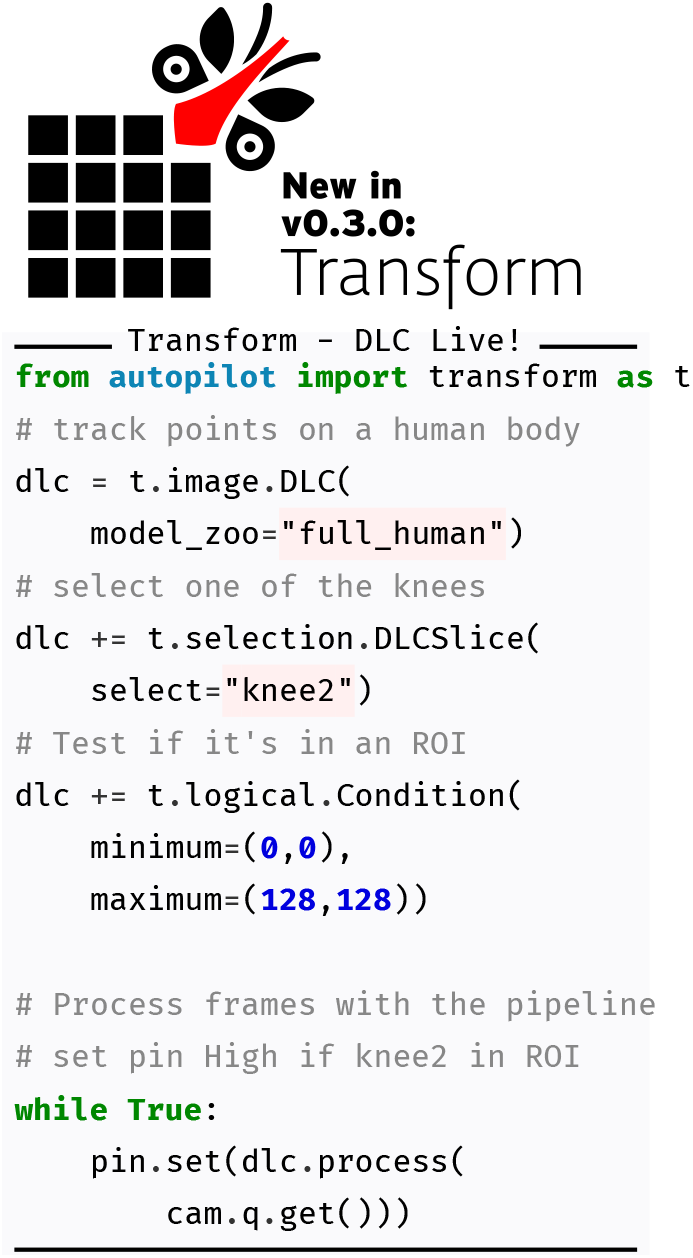
Transforms can be chained together (here with the in-place addition operator `+=`) to make pipelines that encapsulate the logical relationship between some input and a desired output. Here `pin` is a Digital_Out object, and `cam` is a [PiCamera](https://docs.auto-pi-lot.com/en/latest/hardware/cameras.html#autopilot.hardware.cameras.PiCamera) with `queue` enabled.

In addition to computing derived values, we use transforms in a few ways, including

- **Bridging Hardware** — Different hardware devices use different data types, units, and scales, so transforms can [rescale](https://docs.auto-pi-lot.com/en/latest/transform/units.html#autopilot.transform.units.Rescale) and convert values to make them compatible.
- **Integrating External Tools** — The number of exciting analytical tools for realtime experiments keep growing, but in practice they can be hard to use together. The transform module gives a scaffolding for writing wrappers around other tools and exposing them to each other in a shared framework, as we did with DeepLabCut-Live[30], making closed-loop pose tracking available to the rest of Autopilot’s ecosystem. We don’t need to rally thousands of independent developers to agree to write their tools in a shared library, instead transforms make wrapping them easy.
- **Extending Objects** — Transforms can be used to augment existing objects and create new ones. For example, a [motion sensor](https://docs.auto-pi-lot.com/en/latest/hardware/i2c.html#autopilot.hardware.i2c.I2C_9DOF) uses the [spheroid](https://docs.auto-pi-lot.com/en/latest/transform/geometry.html#autopilot.transform.geometry.Spheroid) transform to calibrate its accelerometer, and the [gammatone filter](https://docs.auto-pi-lot.com/en/latest/transform/timeseries.html#autopilot.transform.timeseries.Gammatone)^5^ extends the [Noise](https://docs.auto-pi-lot.com/en/latest/stim/sound/sounds.html#autopilot.stim.sound.sounds.Noise) sound to make a gammatone [filtered noise](https://docs.auto-pi-lot.com/en/latest/stim/sound/sounds.html#autopilot.stim.sound.sounds.Gammatone) sound.

Like Tasks and Hardware, the transform module provides a scaffolding for writing reference implementations of algorithms commonly needed for realtime behavioral experiments. For example, neuroscientists often want to quickly measure a research subject’s velocity or orientation, which is possible with inexpensive inertial motion sensors (IMUs), but since anything worth measuring will be swinging the sensor around with wild abandon the readings first need to be rotated back to a geocentric coordinate frame. Since the readings from an accelerometer are noisy, we found a few whitepapers describing using a Kalman filter for fusing the accelerometer and gyroscope data for a more accurate orientation estimate ([49, 50]), but couldn’t find an implementation. We [wrote one](https://docs.auto-pi-lot.com/en/latest/_modules/autopilot/transform/geometry.html#IMU_Orientation) and integrated it into the IMU class (Figure 3.11). Since it’s an independent transform, it’s available to anyone even if they use nothing else from Autopilot.

**Figure 3.11:**
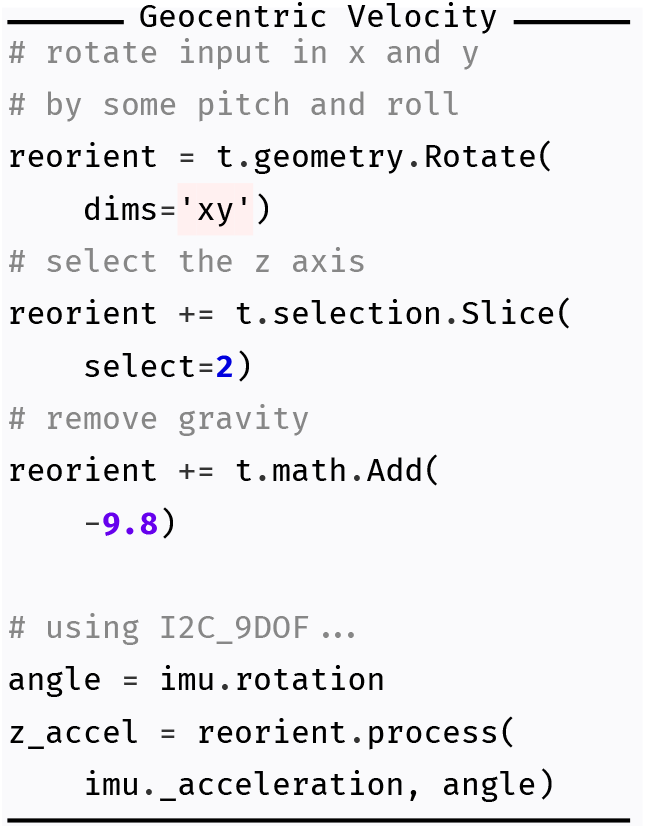
Using the IMU_Orientation transform built into the IMU’s `rotation` property, a processing chain to reorient the accelerometer reading and subtract gravity for geocentric z-axis acceleration.

Transforms were made to be composed, so we broke it into independent sub-operations: A [Kalman](https://docs.auto-pi-lot.com/en/latest/transform/timeseries.html#autopilot.transform.timeseries.Kalman) filter, [rotation](https://docs.auto-pi-lot.com/en/latest/transform/geometry.html#autopilot.transform.geometry.Rotate), and a [spheroid correction](https://docs.auto-pi-lot.com/en/latest/transform/geometry.html#autopilot.transform.geometry.Spheroid) to calibrate accelerometers. Then we combined it with the DLC-Live transform for a fast but accurate motion estimate from position, velocity, and acceleration measurements from three independent sensors. Since each step of the transformation is exposed in a clean API, it was straightforward to [extend the Kalman filter](https://github.com/auto-pi-lot/autopilot-plugin-parallax/blob/759dbb382b90a99f71edf5070772ac18555b67dd/kalman_position.py) to accomodate the the wildly different sampling rates of the camera and IMU. It’s still got its quirks, but that’s the purpose of plugins — to make the code [available and documented](https://wiki.auto-pi-lot.com/index.php/Plugin:Parallax) without formally integrating it in the library.

### 3.6 Stimuli

A hardware object would control a speaker, whereas stimulus objects are the individual sounds that the speaker would play. Like tasks and hardware, Autopilot makes stimulus generation portable between users, and is released with a family of common sounds like tones, noises, and sounds from files. The logic of sound presentation is contained in an inherited metaclass, so to program a new stimulus a user only needs to describe how to generate it from its parameters (Figure 3.12). Sound stimuli are better developed than visual stimuli as of v0.5.0, but we present a proof-of-concept visual experiment (Section 4.5) using [psychopy](https://www.psychopy.org/)[20].

**Figure 3.12:**
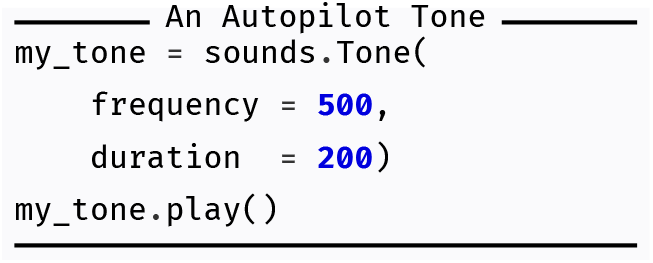
Autopilot stimuli are parametrically defined and inherit all the playback logic that makes them easy to integrate in tasks

Autopilot controls the realtime audio server [jack](http://jackaudio.org/) from an independent Python process that dumps samples directly into jack’s buffer (Figure 3.13), giving it a trigger-to-playback latency very near the theoretical minimum (Section 4.2). Sounds can be pre-buffered in memory or synthesized on demand to play continuous sounds. Because the realtime server is independent from the logic of sound synthesis and storage, stimuli can be controlled independently from different threads without interrupting audio or dropping frames.

**Figure 3.13:**
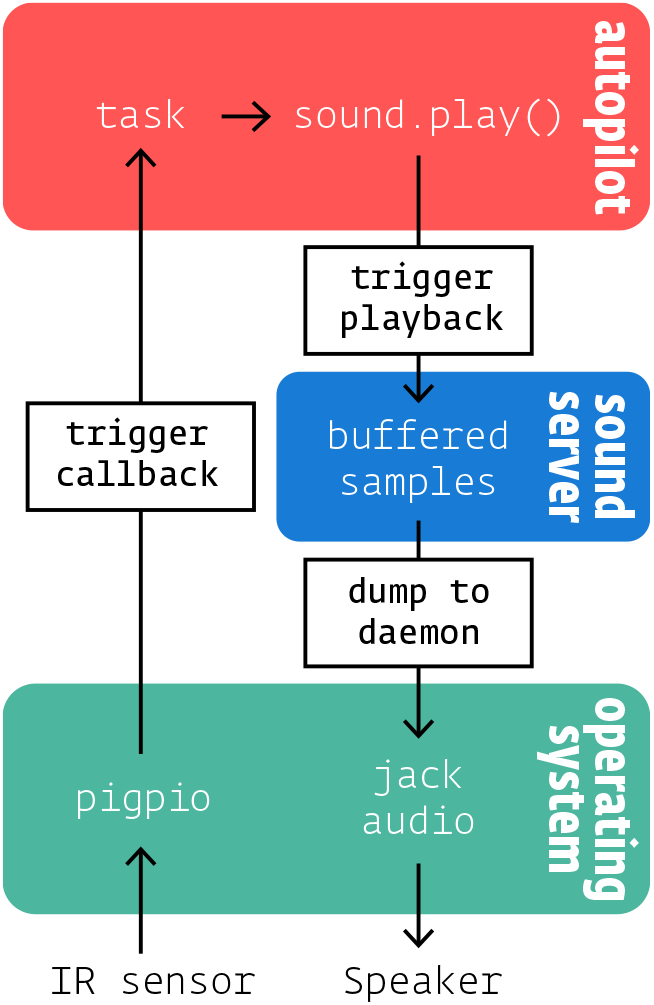
Our sound server keeps audio samples buffered until a `.play()` method is called, and then dumps them directly into the jack audio daemon.

We use the [Hifiberry Amp 2](https://www.hifiberry.com/shop/boards/hifiberry-amp2/), a combined sound card and amplifier, which is capable of 192kHz/24Bit audio playback. Autopilot and Jack can output to any sound hardware, however, including the builtin audio of the Raspberry Pi if fidelity isn’t important. There are no external video cards for the Raspberry Pi 4b^6^, but its embedded video card is capable of presenting video and visual stimuli (Section 4.5) especially if the other computationally demanding parts of the task are distributed to other Raspberry Pis (Section 3.7). If greater video performance is needed, Autopilot is capable of running on typical desktops as well as other single-board computers with GPUs (as we did with the Nvidia Jetson in [30]).

### 3.7 Agents

All of Autopilot’s components can be organized into a single system as an “agent,” the executable that coordinates everyday use. An agent encapsulates:

- **Runtime Logic** — an initialization routine that starts any needed system processes and any subsequent operations that define the behavior of the agent.
- **Networking Station** — Agents have networking objects called `Stations` that are intended to be the “load bearing” networking objects (described more below).
- **Callbacks** — An action vocabulary that maps different types of messages to methods for handling them. Called `listens` to disambiguate from other types of callbacks.
- **Dependencies** — Required packages, libraries, and system reconfigurations needed to operate. Python dependencies are currently defined for agents as groups of [optional packages](https://peps.python.org/pep-0621/#dependencies-optional-dependencies)^7^, and system configuration is done with [scripts](https://docs.auto-pi-lot.com/en/latest/setup/scripts.html) which shorthand common operations like [compiling OpenCV](https://github.com/auto-pi-lot/autopilot/blob/90956187d4222f16f67ab8b39b8359da954d5dcc/autopilot/setup/scripts.py#L140-L183) with optimizations for the raspi or [enabling a soundcard](https://github.com/auto-pi-lot/autopilot/blob/90956187d4222f16f67ab8b39b8359da954d5dcc/autopilot/setup/scripts.py#L92-L100).

Together, these define an agent’s *role* in the swarm.

There are currently two agents in Autopilot:

- **Terminal** - The user-facing control agent.
- **Pilot** - A Raspberry Pi that runs tasks, coordinates hardware, and optionally coordinates a set of child Pis.

**Terminal** agents serve as a root node (see Section 3.8) in an Autopilot swarm. The terminal is the only agent with a GUI, which is used to control its connected pilots and visualize incoming task data. The terminal also manages data and keeps a registry of all active experimental subjects. The terminal is intended to make the day-to-day use of an Autopilot swarm manageable, even for those without programming experience. The terminal GUI is described further in Section 3.9.

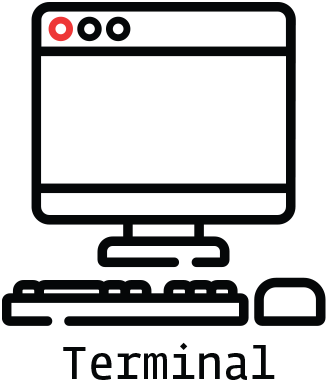

**Pilot** agents are the workhorses of Autopilot—the agents that run the experiments. Pilots are intended to operate as always-on, continuously running system services. Pilots make a network connection to a terminal and wait for further instructions. They maintain the system-level software used for interfacing with the hardware connected to the Raspberry Pi, receive and execute tasks, and continually return data to the terminal for storage.

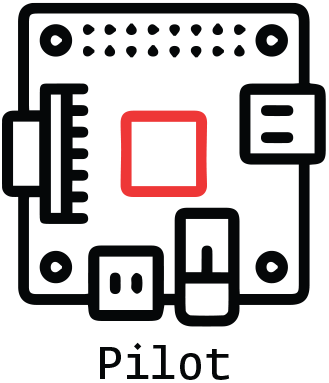

Each agent runs autonomously, and so a Pilot can run a task without a Terminal and store data locally, a Terminal can be used without Pilots to define protocols and manage subjects, and so on. This decoupling lets each agent have more freedom in its behavior at the expense of the complexity of configuring and maintaining them (see Sec. 5.10 and 5.13). All interaction is based on the “listen” callbacks known by the agents, so to start a task a Terminal will send a Pilot a `“START”` message containing information about a `Task` class that it is to run along with its parameterization. The Pilot then attempts to run the task, sends a message to the Terminal alerting it to a `“STATE”` change, and begins streaming data back to it in messages with a `“DATA”` key.

Each pilot is capable of mutually coordinating with one or many **Copilots**^8^. We are still experimenting with, and thus openminded to the best way to structure multi-pilot tasks. Like many things in Autopilot, there is no one right way to do it, and the strategy depends on the particular constraints of the task. We include a few examples in the network latency and go/no-go tasks in the [plugin](https://wiki.auto-pi-lot.com/index.php/Plugin:Autopilot_Paper) that accompanies this paper, and expand on this a bit further in a few parts of section 5, as it is a major point of active development.

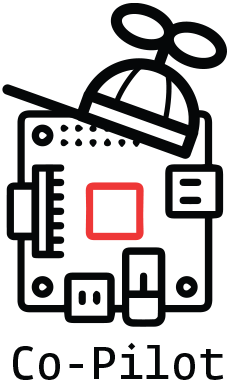

#### Behavioral topologies

We think one of the most transformative features of Autopilot’s distributed structure is the control that users have over what we call “behavioral topology.” The logic of hardware and task operation within an agent, the distribution of labor between agents performing a task, and the pattern of connectivity and command within a swarm of agents constitute a topology.

Below we illustrate this idea with a few examples:

- **Pilot Swarm** - The first and most obvious topological departure from traditional behavioral instrumentation is the use of a single computer to independently coordinate tasks in parallel. Our primary installation of Autopilot is a cluster of10 behavior boxes that can independently run tasks dispatched from a central terminal which manages data and visualization. This topology highlights the expandability of an Autopilot system: adding new pilots is inexpensive, and the single central terminal makes controlling experiments and managing data simple.
- **Shared Task** - Tasks can be shared across a set of copilots to handle tasks with computationally intensive operations. For example, in an open-field navigation task, one pilot can deliver position-dependent sounds while another records and analyzes video of the arena to track the animal’s position. The terminal only needs to be configured to connect to the parent pilot, but the other copilot is free to send data to the Terminal marked for storage in the subject’s file as well.
- **Distributed Task** - Many pilots with overlapping responsibilities can cooperate to perform distributed tasks. We anticipate this will be useful when the experimental arenas can’t be fully contained (such as natural environments), or when experiments require simultaneous input and output from multiple subjects. Distributed tasks can take advantage of the Pi’s wireless communication, enabling, for example, experiments that require many networked cameras to observe an area, or experiments that use the Pis themselves as an interface in a multisubject augmented reality experiment.
- **Multi-Agent Task** - Neuroscientific research often consists of multiple mutually interdependent experiments, each with radically different instrumentation. Autopilot provides a framework to unify these experiments by allowing users to rewrite core functionality of the program while maintaining integration between its components. For example, a neuroethologist could build a new “Observer” agent that continually monitors an animal’s natural behavior in its home cage to calibrate a parameter in a task run by a pilot. If they wanted to manipulate the behavior, they could build a “Compute” agent that processes Calcium imaging data taken while the animal performs the task to generate and administer patterns of optogenetic stimulation. Accordingly, passively observed data can be combined with multiple experimental datasets from across the subject’s lifespan. We think that unifying diverse experimental data streams with interoperable frameworks is the best way to perform experiments that measure natural behavior in the fullness of its complexity in order to understand the naturally behaving brain[25].

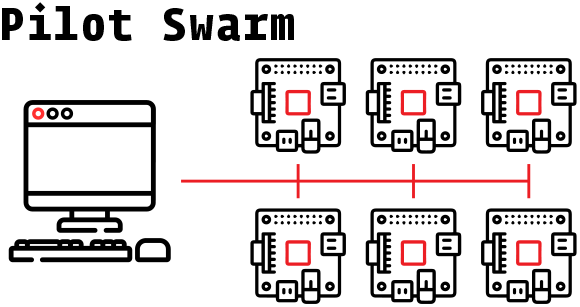

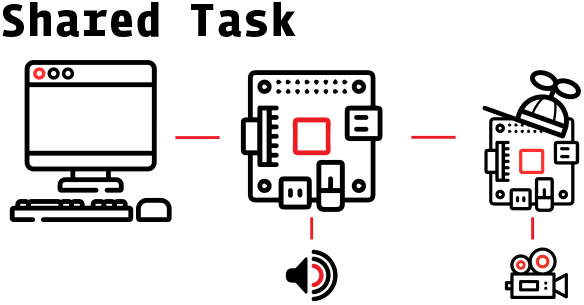

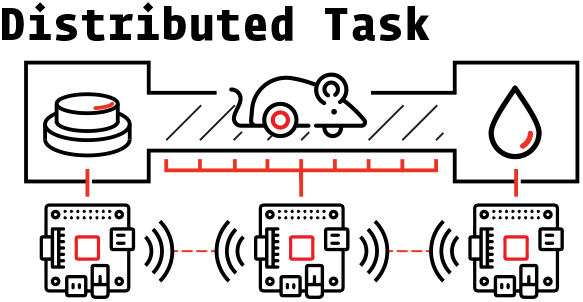

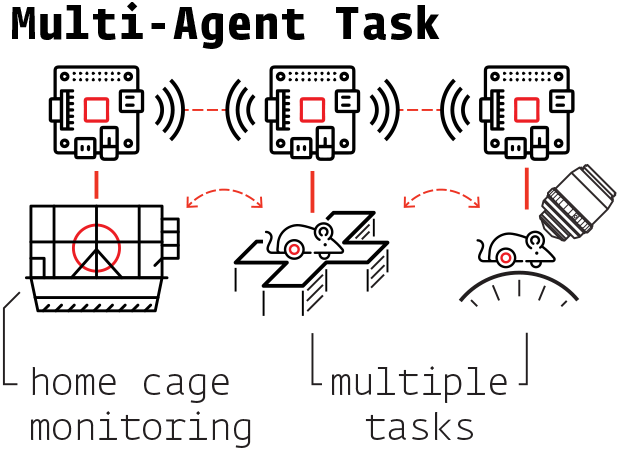

### 3.8 Networking

Agents use two types of object to communicate with one another: core **station** objects and peripheral **node** objects (Figure 3.14). Each agent creates one station in a separate process that handles all communication *between* agents. Stations are capable of forwarding data and maintaining agent state so the agent process is not unnecessarily interrupted. Nodes are created by individual modules run within an agent—eg. tasks, plots, hardware—that allow them to send and receive messages within an agent, or make connections directly to other nodes on other agents after the station discovers their network addresses. Messages are TCP packets^9^, so there is no distinction between sending messages within a computer, a local network, or over the internet^10^.

**Figure 3.14:**
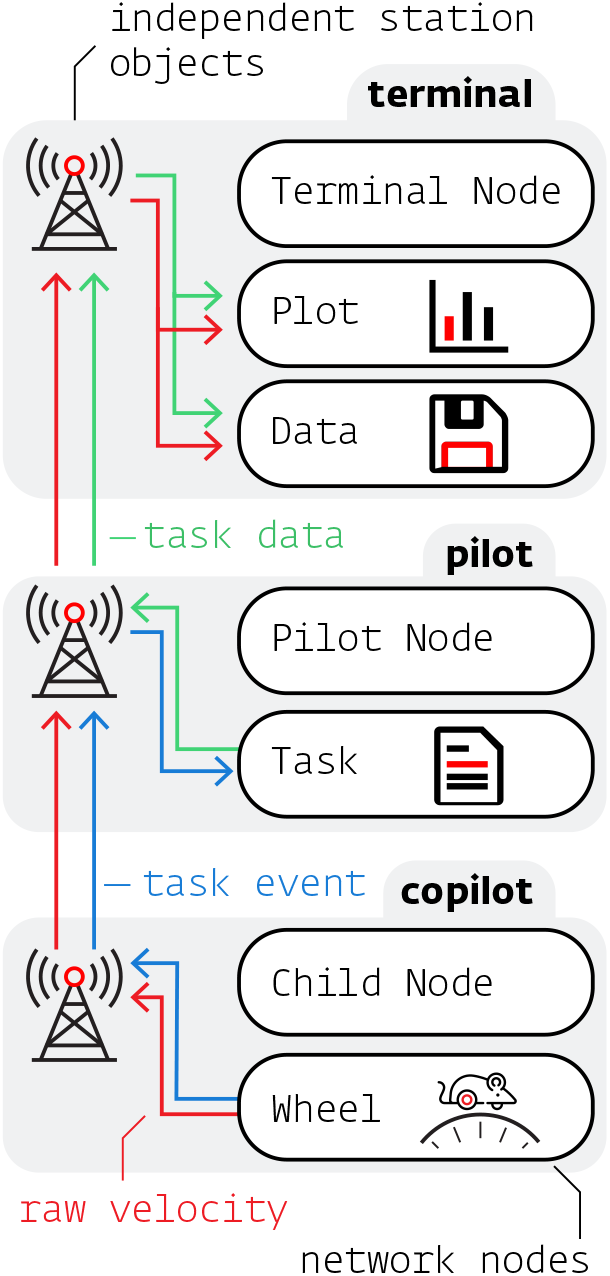
Autopilot segregates data streams efficiently—eg. raw velocity (red) can be plotted and saved by the terminal while only the task-relevant events (blue) are sent to the primary pilot. The pilot then sends trial-summarized data to the terminal (green).

Both types of networking objects are tailored to their hosts by a set of callback functions — **listens** — that define how to handle each type of message. Messages have a uniform key-value structure, where the key indicates the listen used to process the message and the value is the message payload. This system makes adding new network-enabled components trivial:

**Listing 7:**
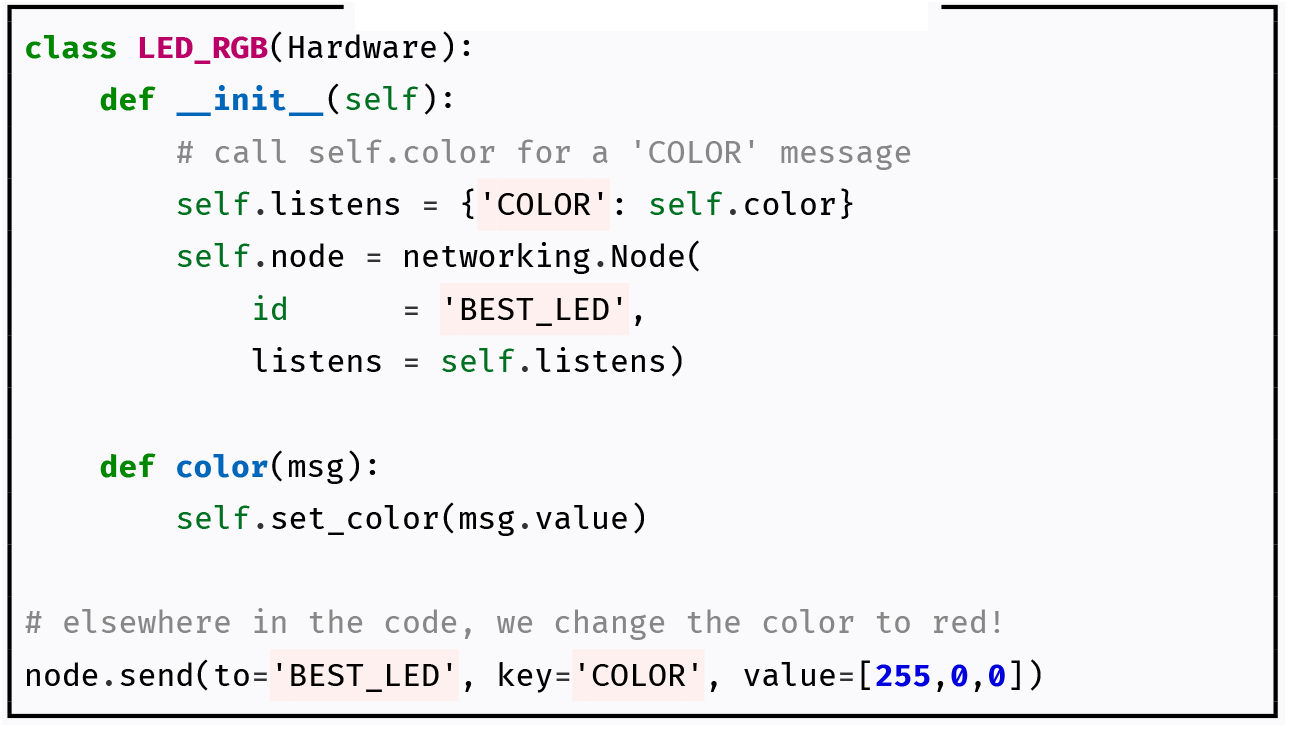
A new networked LED.

Messages are serialized^11^ with JSON, and can handle arrays, including on-the-fly compression with [blosc](https://www.blosc.org/). Net Nodes can create additional sockets to stream data that is stashed in a [queue](https://docs.python.org/3/library/queue.html#queue.Queue), and can take advantage of message batching and compressing multiple arrays together when latency is less critical.

Network connectivity is currently treelike by default (Figure 3.15) — each independent networking object can have many children but at most one parent. This structure makes an implicit assumption about the anisotropy of information flow: ‘higher’ nodes don’t need to send messages to the ‘lowest’ nodes, and the ‘lowest’ nodes send all their messages to one or a few ‘higher’ nodes. It enforces simplified delegation of responsibilities in both directions: a terminal shouldn’t need to know about every hardware object connected to all of its connected pilots, it just sends messages to the pilots, who handle it from there. A far-downstream node shouldn’t need to know exactly how to send its data back to the terminal, so it pushes it upstream until it reaches a node that does.

**Figure 3.15:**
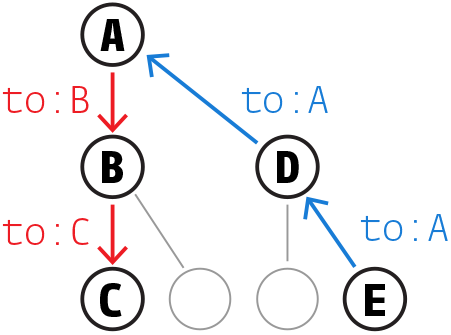
Treelike network structure—downstream messages are addressed by successive nodes, but upstream messages can always be pushed until the target is found.

**Figure 3.16:**
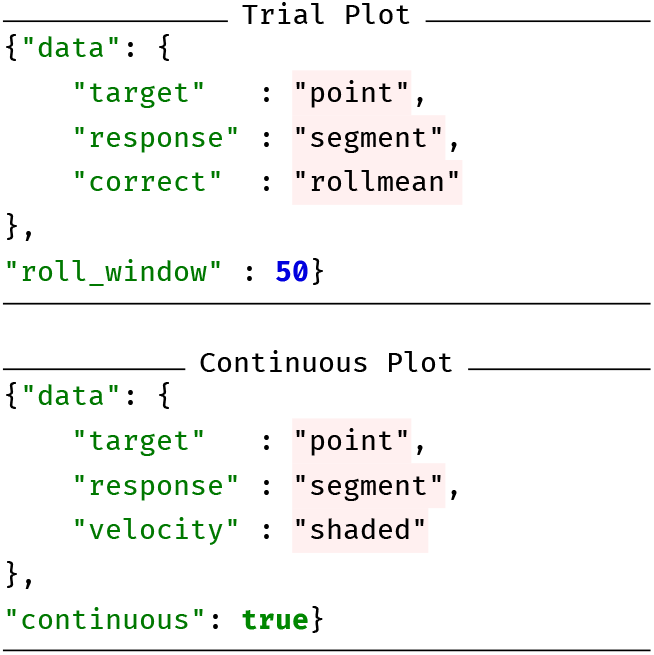
`PLOT` parameters for Figure 3.17. In both, “target” and “response” data are mapped to “point” and “segment” graphical primitives, but timestamps rather than trial numbers are used for the x-axis in the “continuous” plot (Figure 3.17, bottom). Additional parameters can be specified, eg. the trial plot (Figure 3.17, top) computes rolling accuracy over the past 50 trials

**Figure 3.17:**
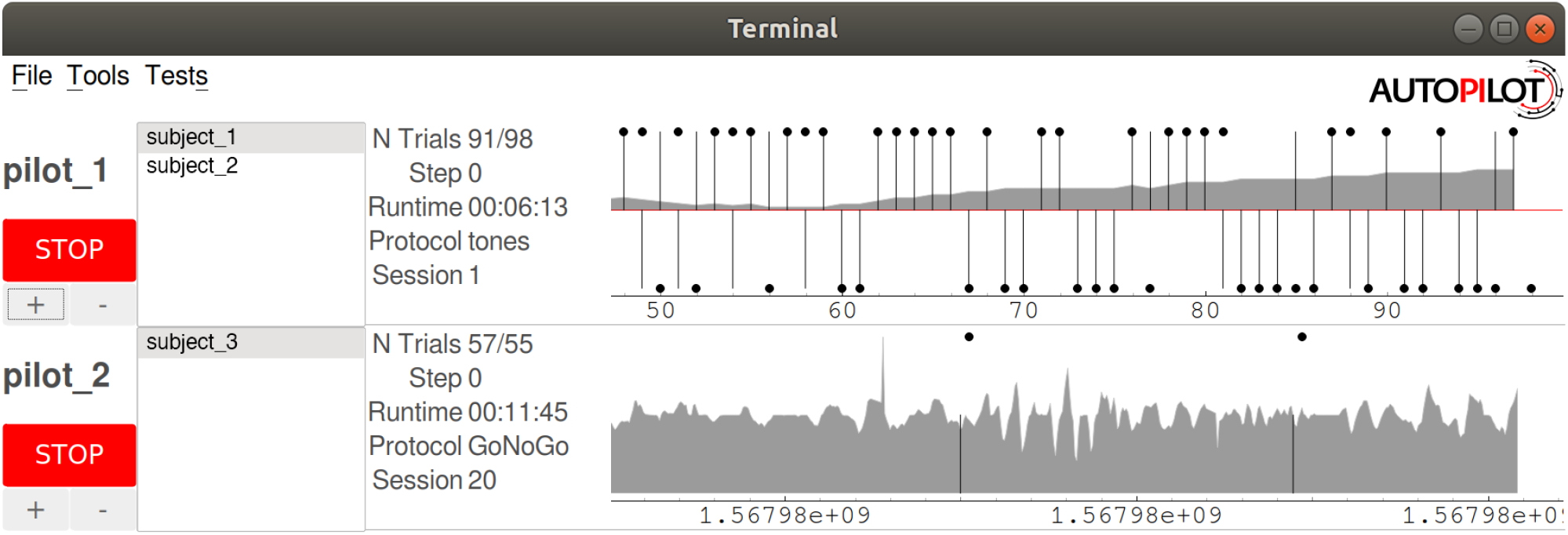
Screenshot from a terminal GUI running two different tasks with different plots concurrently. `pilot_1` runs 2 subjects: (`subject_1` and `subject_2`), while `pilot_2` runs `subject_3`. See Figure 3.16 for plot description

This treelike structure is useful for getting started quickly with the default full system configuration, but for experimenting with different configurations it is also possible to directly connect network nodes. The Station backbone is then a useful way of connecting objects across agents, as messages can be sent as multihop messages through a connected tree of stations^12^ to make an initial connection without hard-coding an IP or Port. In the future we plan to simplify this further by directly implementing a peer to peer discovery model, see Section 5.6.

### 3.9 GUI & Plots

The terminal’s GUI controls day-to-day system operation^13^. It is intended to be a nontechnical frontend that can be used by those without programming experience.

For each pilot, the terminal creates a control panel that manages subjects, task operation, and plots incoming data. [Subjects can be managed](http://docs.auto-pi-lot.com/guide.training.html#creating-a-subject) through the GUI, including creation, protocol assignment, and metadata editing. Protocols can also be [created from within the GUI](http://docs.auto-pi-lot.com/guide.training.html#creating-a-protocol). The GUI also has a set of basic maintenance and informational routines in its menus, like calibrating water ports or viewing a history of subject weights.

The simple callback design and network infrastructure makes adding new GUI functionality straightforward, and in the future we intend to extend the plugin system such that plugins can provide additional menu actions, plots, and utilities.

#### Plotting

Realtime data visualization is critical for monitoring training progress and ensuring that the task is working correctly, but each task has different requirements for visualization. A task that has a subject continuously running on a ball might require a continuous readout of running velocity, whereas a trial-based task only needs to show correct/incorrect responses as they happen. Autopilot approaches this problem by assigning the data returned by the task to graphical primitives like points, lines, or shaded areas as specified in a task’s `PLOT` dictionary (taking inspiration from Wilkinson’s grammar of graphics[51]).

The GUI is now some of the oldest code in the library, and we are in the process of decoupling some of its functionality from its visual representation and moving to a model where it is a thinner wrapper around the data modeling tools. Following the lead of formal models with strict typing will, for example, make plotting more fluid where the researcher can map incoming data to the set of graphical elements that are appropriate for its type. We discuss this further in section 5.8

## 4 Tests

We have been testing and refining Autopilot since we built our swarm of 10 training boxes in the spring of 2019. In that time 178 mice^1^ have per-formed over 6 million trials on a range of tasks. While Autopilot is still relatively new, it is by no means untested.

In this section we will present a set of basic performance benchmarks while also showing several of the different ways that Autopilot can be used. The code for all of the following tests is available as a [plugin](https://github.com/auto-pi-lot/plugin-paper) that is further documented on the [wiki](https://wiki.auto-pi-lot.com/index.php/Plugin:Autopilot_Paper), and runs on a prerelease of v0.5.0. Materials tables (Table 4.1) for each test link more specifically to the test code and provide additional hardware and version documentation, where appropriate.

**Table 4.1:** General Materials

| Hardware |  |
| --- | --- |
| Raspi | Raspberry Pi 4b |
| Oscilloscope | Rigol DS1054Z |
| Software |  |
| Autopilot | v0.5.0a |
| Plugin | Autopilot_Paper |
| Python | 3.9.12 |
| RaspiOS | Bullseye 22-04-04 (lite) |
| Analysis |  |
| R | 4.2.0 |
| ggplot2[52] | 3.3.5 |
| dplyr[53] | 1.0.9 |
| purrr[54] | 0.3.4 |
| pandas[45] | 1.4.2 |
| numpy[23, 55] | 1.21.6 |

### 4.1 GPIO Latency

Neurons compute at millisecond timescales, so any task that links neural computation to behavior needs to have near-millisecond latency. We start by characterizing Autopilot’s GPIO control latency in “script mode” — using the GPIO control classes on their own, without using any of the rest of Autopilot’s modules.

#### Output Latency

We first tested the software measured latency between when a command to write a value to a GPIO pin is issued and when it completes (Figure 4.1, Table 4.2). By default, the pigpio interface we use to control GPIO pins issues a command and then confirms the request was successful by querying the pigpio daemon for the status of the pin (Write+Read). We [extended](https://github.com/sneakers-the-rat/pigpio/commit/0782de06f0a5c092063118733ad2df9d65f1f1a0) pigpio to just issue the command without confirmation to estimate the true time between when the command is issued and when the voltage of the pin changes (Write). Each of these operations takes roughly 40*μs* with minimal jitter (Median ± IQR — Write Only: 40.8*μs*±0.41, Write and Read: 88.4 *μs*±2.02, n=100,000 each).

**Figure 4.1:**
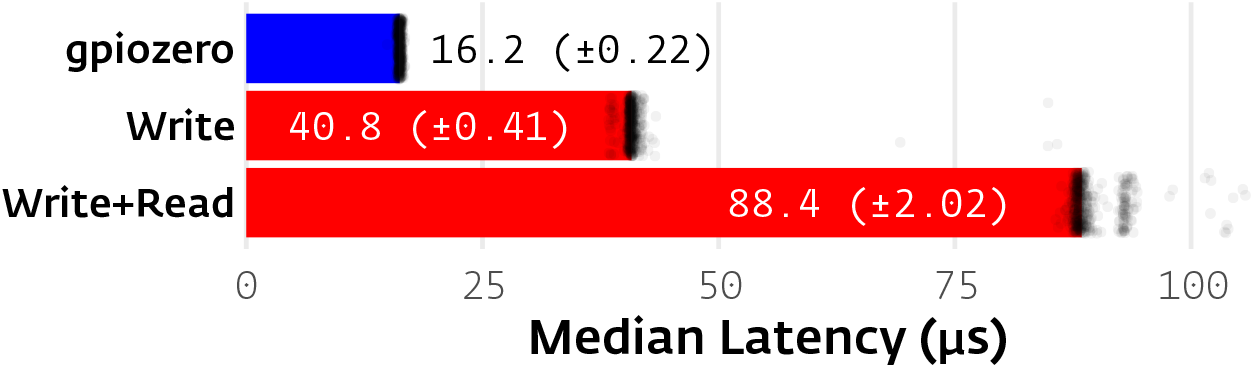
Software latency from GPIO write to completion of command. Values are presented as medians ± IQR with n=100,000 tests for each. A random subsample of 500 (for tractability of plotting) of each type of test are presented (black points) after filtering to the bottom 99th percentile to exclude extreme outliers. Commands sent using pigpio (red) took roughly 40*μs* each (write and read are effectively two separate commands, Write Only: 40.8*μs*±0.41, Write and Read: 88.4 *μs*±2.02). The prototype gpiozero wrapper (blue) using RPi.GPIO as its backend was faster, taking 16.2*μs*±0.22 to complete.

**Table 4.2:** GPIO Latency Materials. (Parameters in {} are input in separate runs)

|  |  |
| --- | --- |
| Code | test_gpio.py |
| replicate | python test_gpio.py -w {0,1,2} -n 100000 |
| gpiozero | 1.6.2 |
| pigpio | 3c23715 |

Pigpio is useful as a general purpose controller because of its ability to run scripts within its daemon, use hardware PWM via direct memory access, and consistently poll for pin state, but takes a latency penalty because the python interface communicates with it through a local TCP socket. To demonstrate the flexibility of Autopilot in incorporating additional software libraries, we wrote a [thin wrapper](https://github.com/auto-pi-lot/plugin-paper/blob/main/plugin_tests/hardware/zero.py) around [gpiozero](https://gpiozero.readthedocs.io/en/stable/), which can use RPi.GPIO to directly write to the GPIO registers. For simple output, this wrapper proved to be faster (16.2 *μs* ± 0.22, n=100,000), and with 63 lines of code is now available in the plugin accompanying this paper to be used, repurposed, and extended.

#### Roundtrip Input/Output Latency

Output commands usually aren’t issued in isolation, but as a response to some external or task-driven trigger. We measured the roundtrip latency from a 5V square pulse from an external function generator to when an output pin was flipped from low to high on an oscilloscope (Table 4.3).

**Table 4.3:**
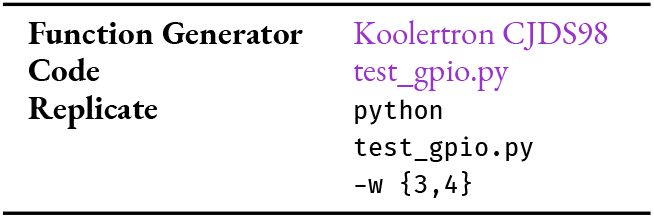
Roundtrip Latency Materials

Typically that is as much methodological detail as you would expect in a scientific paper, but actually making those measurements via oscilloscope requires knowing how to set up such a test as well as how to extract the measured data afterwards — which is not altogether trivially available technical knowledge. As an example of how integrating semantically linked documentation with experimental tools enables a fundamentally deeper kind of reproducibility and methodological transparency, we instead documented these operations, including a code sample and a guide to unlocking additional features on our oscilloscope on the [autopilot wiki](https://wiki.auto-pi-lot.com/index.php/Rigol_DS1054Z). The code to extract traces from the oscilloscope is also included in this paper’s [plugin](https://wiki.auto-pi-lot.com/index.php/Plugin:Autopilot_Paper), which links to the oscilloscope page with a `[[Controls Hardware::Rigol DS1054Z]]` tag, so it is possible to bidirectionally find code examples from the oscilloscope page as well as find further documentation about the hardware used in this paper from the plugin page. The same can be true for any hardware used by any plugin in any paper using Autopilot.

We used the `assign_cb` method of the Digital_In class to test the typical roundtrip latency that Autopilot objects can deliver. This gave us a median 474 *μs* (IQR: 52.5*μs*) latency (Red in Figure 4.3). GPIO callbacks are flexible, and can use arbitrary python functions, but if all that’s needed is to trigger one pin off of another with some simple logic like a parametric digital waveform or static “on” time, pigpio also allows us to directly program pin to pin logic as a “[pigs](http://abyz.me.uk/rpi/pigpio/pigs.html)” script (literally Figure 4.2) that runs within the pigpio daemon. The pigs script gave us roughly three orders of magnitude lower latency (Median ± IQR: 370ns ± 140, blue in 4.3).

**Figure 4.2:**
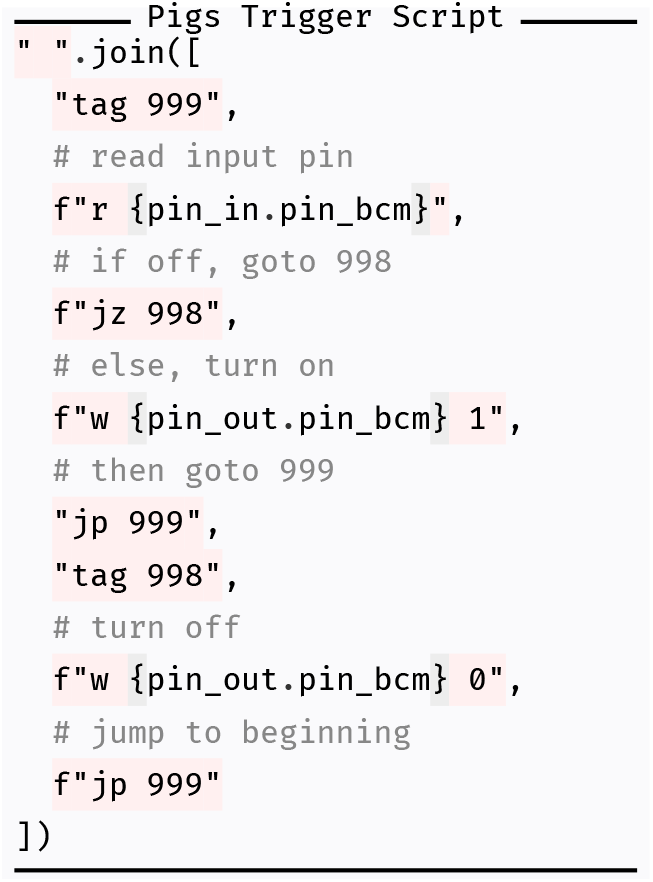
The pigs script used to trigger one pin (pin_out), from another (pin_in). At the expense of a little bit of complexity having to write a script in its scripting language, we are able to reduce latency by three orders of magnitude.

**Figure 4.3:**
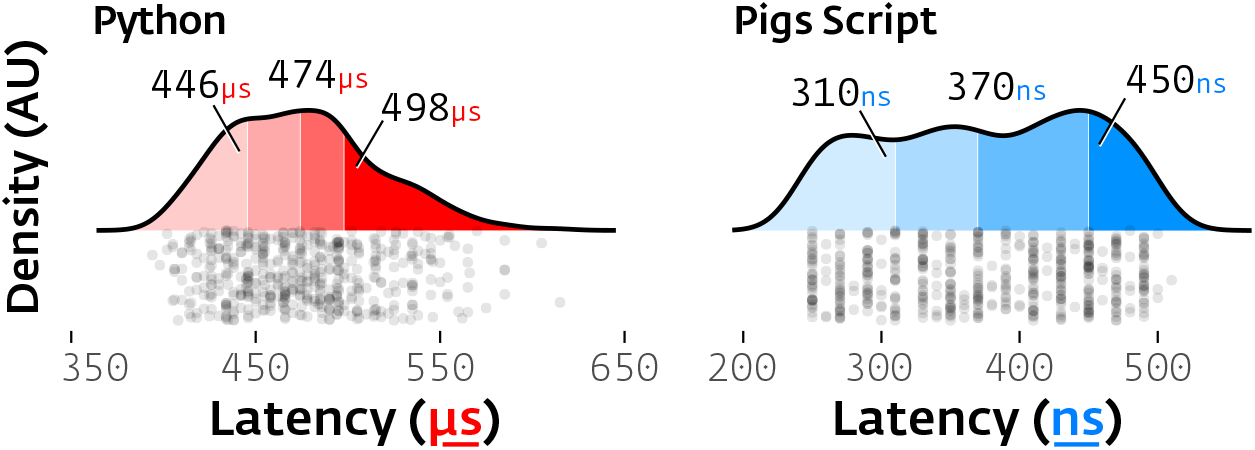
Roundtrip latency from external trigger to digital output using two methods: Typical Autopilot callback function given to a `Digital_In` object that turns a `Digital_Out` pin on for 1ms when an input pin changes state (Red, Left, Median ± IQR: 474 *μs*± 52.5). Pigs script that runs entirely within the pigpio daemon (Blue, Right, 380ns ± 140). For each, black points represent individual measurements (n=525), annotations are quartiles.

### 4.2 Sound Latency

We measured end to end, hardware input to sound output latency by measuring the delay between an external digital input and the onset of a 10kHz pure tone (Table 4.4). Sound playback was again triggered by the `Digital_In` class’s callback method, and sound samples were buffered in a [deque](https://docs.python.org/3/library/collections.html#collections.deque) held in a separate process by the jack audio client between each trial. A `Digital_Out` pin was wired to the `Digital_In` pin in order to deliver the trigger pulse (but the `Digital_Out` pin was uninvolved in the software trigger for sound output).

**Table 4.4:** Sound Latency Materials

|  |  |
| --- | --- |
| **Sound Card** | Hifiberry Amp2 |
| **jack** | 1.9.22 |
| **Code** | test_sound.py |
| **Replicate** | python test_sound.py |

<sup>2</sup> A previous version of this paper included benchmarking and comparison to Bpod and pyControl’s sound onset latency, but since then both packages have changed substantially, including Bpod creating a new `hifi sound module` based off HiFiBerry hardware very similar to the card used here, making those benchmarks obsolete. In this version we have omitted comparative benchmarks in favor of allowing the maintainers of those packages to publish their own benchmarks.

Autopilot’s [jack](http://jackaudio.org/) audio backend was configured with a `192kHz` sampling rate with a buffer with two periods of of 32 samples each for theoretical minimum latency of `0.33ms`^2^. We observed a median 1.35ms (± 0.72 IQR) latency across 521 samples — roughly 4x the theoretical minimum (Figure 4.4). This suggests that Autopilot eliminates most perceptible end-to-end latency, which is necessary for tasks that require realtime feedback. One clear future direction is to write the sound processing loop in a compiled language exposed with a foreign function interface (FFI) to decrease both latency and jitter.

**Figure 4.4:**
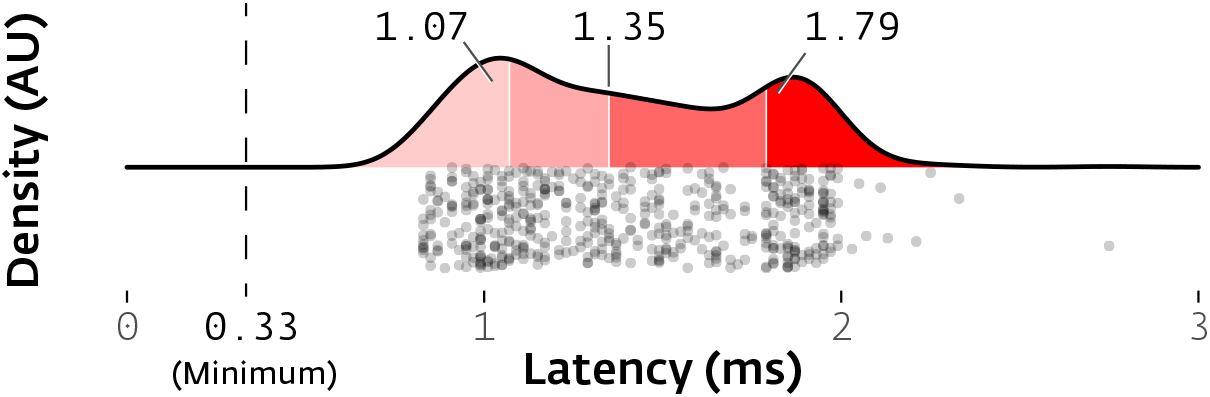
Autopilot has a median 1.35ms (±.72 IQR) latency between an external trigger and sound onset. Individual trials (dots, n=521) are shown beneath a density plot (red area under curve) colored by quartile (shades, numbers above are median, first, and third quartile). This latency is roughly 4x the theoretical minimum (0.33ms, dashed line).

### 4.3 Network Latency

To support data-intensive tasks like those that require online processing of video or electrophysiological data, the networking modules at the core of Autopilot need high bandwidth and low latency.

To test the latency of Autopilot’s networking modules, we switch from “script mode” to “Task mode” (Table 4.5). Tasks are useful for encapsulating multistage routines across multiple devices that would be hard to coordinate with scripts alone. Our `Network_Latency` task consists of one “leader” pilot sending timestamped messages to a “follower” pilot which returns the timestamp marking when it received the message. The two pis communicate via two directly connected `Net_Nodes` (rather than routing each message through agent-level `Station` objects) after the leader pi initiates the follower with a multihop “START” message routed through a Terminal agent containing the task and networking parameters. We measured latency using software timestamps while synchronizing the clocks of the two pis with Chrony, an NTP daemon previously measured to synchronize Raspberry Pis within dozens of microseconds[56]^3^, with the leader pi hosting an NTP server and the follower pi synchronizing its clock solely from the leader. We documented this [on the wiki](https://wiki.auto-pi-lot.com/index.php/NTP) too, since synchronization is a universal problem in multi-computer experiments.

**Table 4.5:** Network Test Materials

|  |  |
| --- | --- |
| **Router** | TP-Link AC1750 |
| **Chrony** | 4.0 |
| **zmq** | 22.3.0 |
| **Code** | Network_Latency |
| **Replicate** | Assign task to subject from Terminal, start Task. |

<sup>3</sup> Our sync is likely to be near to or better than that reported in [56]: in addition to a quiet network, we configured chrony to poll more frequently and tolerate a smaller error than default

Point to point latency was 0.975ms (median, ± 0.1 IQR, n=10,000, Figure 4.5) with some clear bimodality where a subset of messages (2,300 of 10,000) took longer (median 1.567ms). The source of the bimodality is unclear to us, though it could be due to network congestion or interruption by other processes as the networking modules are not run in their own process like the sound server. This latency includes message serialization and deserialization by the builtin JSON library, which is on the order of roughly 100*μs* each for even the very small messages sent in this test. In future versions we will explore other serialization tools like [msgpack](https://msgpack.org/) and offer them as alternate serialization backends.

**Figure 4.5:**
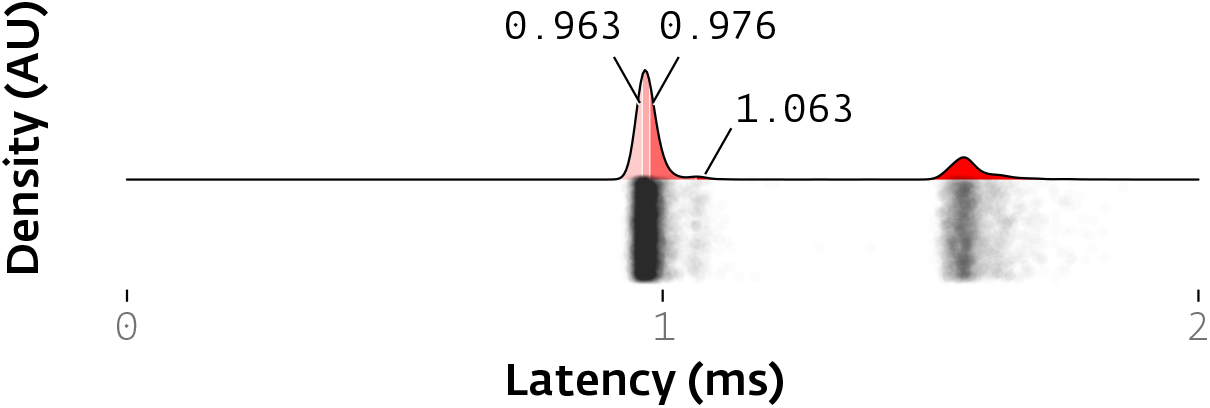
Network latency from when a message is sent from one pilot to when it is received by another. Messages took 0.975ms to send and receive (median, ± 0.1 IQR, n=10,000, overlaid numbers and red shading in density plot indicate quartiles). There is a clear bimodality in latencies for individual messages (black dots, jittered in y-axis) with unclear cause.

### 4.4 Network Bandwidth

To test Autopilot’s bandwidth, we demonstrate yet another modality of use, using Autopilot’s Bandwidth_Test widget, an action available from the Terminal GUI’s `tests` menu that corresponds to a callback “listen” method in the Pilot (Table 4.6). This test requests that one or several pilots send messages at a range of selected frequencies and payload sizes back to the terminal. The messages pass through four networking objects en route: the stations and network nodes running the test for both the terminal and pilots (See Figure 3.14).

**Table 4.6:** Bandwidth Test Materials.

|  |  |
| --- | --- |
| **Terminal** | Macbook Pro 2019, macOS 12.3.1, 2.4 GHz 8-Core Intel Core i9 |
| **blosc2** | v0.2.0 |
| **Code** | `Bandwidth_Test`, `Pilot.l_bandwidth` |
| **Replicate** | Terminal > Tools > Test Bandwidth |

<sup>4</sup> $(1920 * 1080 * 3 * 8 \text{ bits}) / 8 = 6$ megabytes per frame of a 1080p color video, which is why video is rarely streamed uncompressed
<sup>5</sup> maximum average rate of 5000 messages for each of the equivalent empty message tests in the four conditions described below
<sup>6</sup> In all cases, float64 numpy arrays encoded in base64

The needs for streaming experimental data vary depending on what is being streamed. Electrophysiological data is an n-electrode length vector sampled at a rate of dozens to hundreds of kilohertz, so each individual message isn’t very large but there are a lot of them. Video data is a width by height (and for color video, by channel) array that can be relatively large^4^, but it is captured at dozens to hundreds of hertz. Different data streams also have different degrees of compressibility: noisy, quasirandom electrical signals compress relatively poorly, while the typical behavioral neuroscientist’s video of an animal that takes up 1/10th of the frame against a white background can have compression ratios in the hundreds.

Autopilot tries to provide flexibility for streaming different data types by offering message batching and optional on-the-fly compression with [blosc](https://www.blosc.org/). The bounds on bandwidth are then the speed at which an array can be compressed and the rate at which messages of a given size can be sent.

Autopilot’s networking modules were able to send an “empty” (402 byte) message with headers describing the test but no payload at a maximum observed rate of 1,818Hz^5^. Approximately 15% of the duration is spent in message serialization, as a “frozen” preserialized message can be sent at 2,100Hz, though we imagine the need to send the same message thousands of times is rare.

We tested four types of messages with nonzero array^6^ payloads: since the entropy of an array determines how compressible it is, we sent random and all-zero arrays with and without compression. The random and all-zero arrays are the floor and ceiling of compressibility, respectively. Compression gives us two notions of bandwidth: the literal number of bytes that can be passed through a connection, and the effective bandwidth of the size of the arrays that can be transferred with a given compression ratio. We refer to these as “message” and “payload” bandwidth, respectively in Figure 4.6. Message bandwidth reflects the hardware limitations of the Raspberry Pi, but payload bandwidth is the number that matters in practice, as it measures the actual “speed of data” that can be used by the receiver.

**Figure 4.6:**
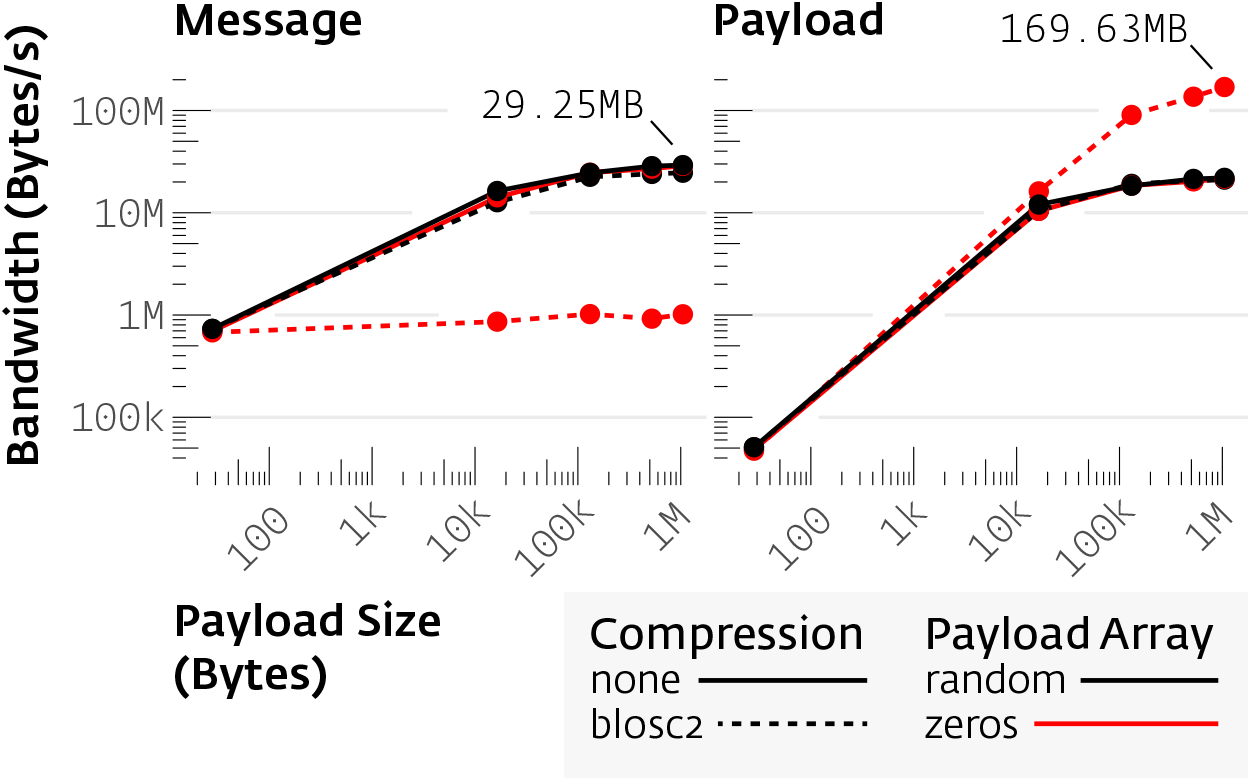
Bandwidth measurements between a pilot and terminal for compressed (solid lines) or uncompressed (dotted lines) arrays of random numbers (black) or zeros (red, each point n=5000 messages). As message size increased, the bandwidth for the rate of bytes transferred in serialized messages (“message bandwidth,” left) plateaued at 29.25MBytes/s, while the effective bandwidth of arrays before and after compression (“payload bandwidth,” right) reached 169.63MBytes/s. Real data will fall somewhere in this effective bandwidth range, depending on its compressibility.

As we increased the size of the array payload^7^, the message bandwidth plateaued at a maximum of 29.25MByte per second (Figure 4.6, left). After this plateau, increasing the message size trades off linearly with the rate of messages sent. For all but the compressed array of zeros, the payload bandwidth mirrored the message bandwith with some trivial overhead from the base64 encoding. The compressed array of zeros, however, had an effective payload bandwidth of 169.6MBytes/s, a compromise between the speed of compression with the smaller message size^8^. The compressed random array had only negligible differences in payload and message bandwidth compared to the uncompressed random array, indicating that the overhead for blosc is trivial.

The ability to batch messages allows researchers to tune the size of an individual message to their particular need for high bandwidth or low latency. Since the compressibility of real data varies across the entire entropic range from randomness to arrays of all zeros, Autopilot doesn’t have a single “bandwidth”, but one that ranges between 30 and 170MByte/s^9^.This bandwidth makes Autopilot capable of streaming raw Calcium imaging^10^ and electrophysiological data from modern high-density probes^11^. Its flexible architecture allows researchers to decide how to build their experiments by distributing different components over different combinations of computers: stream data from a raspberry pi to a more powerful computer for processing, use GPIO rather than network triggers for time-critical operations — meeting the tooling challenge of complex, hardware-intensive, multimodal experiments that define contemporary systems neuroscience.

### 4.5 Distributed Go/No-go Task

We designed a visual go/no-go task as a proof of concept for distributing task elements across multiple Pis, and also for the presentation of visual stimuli (Figure 4.7, Table 4.7). While the rest of the tests presented have been re-run, in the time since the intitial publication of the preprint we have not done substantial work on Autopilot’s visual stimulus module, and so this section is presented as previously written using the v0.1.0 initial release.

**Figure 4.7:**
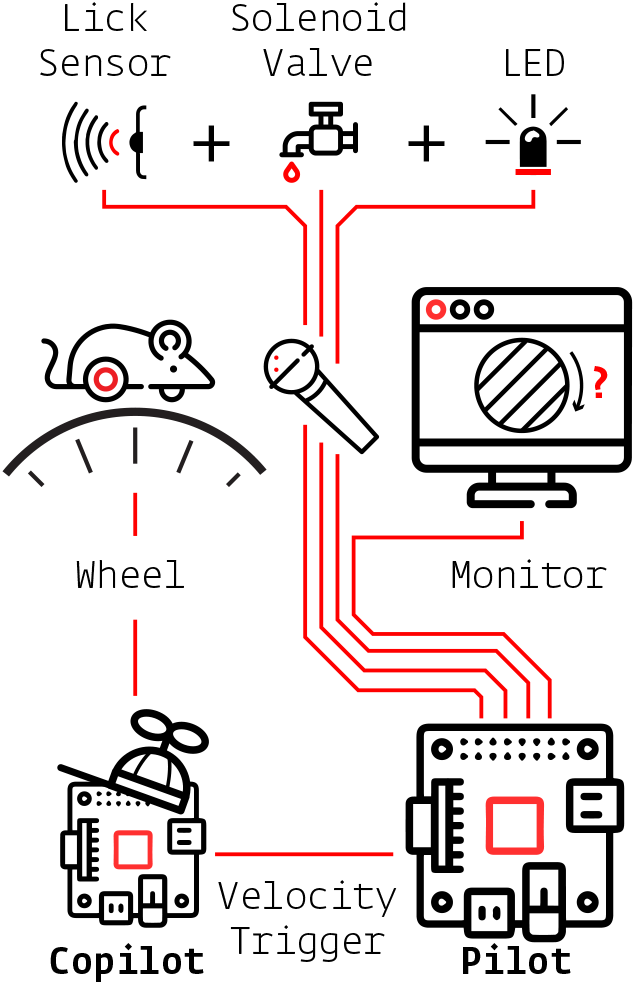
Hardware distribution for the distributed go/no-go task. Red lines indicate physical connections between hardware components. The lick sensor, solenoid valve, and LED are physically bundled into one component represented as the mouse’s microphone.

**Table 4.7:** Go/No-go Materials

|  |  |
| --- | --- |
| **Hardware** |  |
| Beam Break | TT Electronics OPB901L55 |
| Monitor | Acer S230HL |
| Lick Port | Autopilot Tripoke v1 |
| Light Sensor | Thorlabs PM100D |
| **Software** |  |
| psychopy | v3.1.5 |
| glfw | v1.8.3 |

In this task, a head-fixed subject would^12^ be running on a wheel in front of a display with a lick-detecting water port able to deliver reward. Above the port is an LED. Whenever the LED is green, if the subject drops below a threshold velocity for a fixation period, a grating stimulus at a random orientation is presented on the monitor. After a random delay, there is a chance that the grating changes orientation by a random amount. If the subject licks the port in trials when the orientation is changed, or refrains from licking when it is not, the subject is rewarded.

One pilot controlled the operation of the task, including the coordination of a copilot. The pilot was connected to the LED and solenoid valve for reward delivery, as well as a monitor^13^ to display the gratings^14^. The copilot continuously streamed velocity data (measured with a USB optical mouse against the surface of the wheel) back to the terminal for storage (see also Figure 3.14, which depicts the network topology for this task). The copilot waited for a message from the pilot to initiate measuring velocity, and when a rolling average of recent velocities fell below a given threshold the copilot sent a TTL trigger back to the pilot to start displaying the grating. This split-pilot topology allows us to poll the subject velocity continuously (at 125Hz in this example) without competing for resources with psychopy’s rendering engine.

We measured trigger (TTL pulse from the copilot) to visual stimulus onset latency using the measurement cursors of our oscilloscope as before. To detect the onset of the visual stimulus, we used a high-speed optical power meter attached to the top-left corner of our display monitor. The stimulus was a drifting Gabor grating drawn to fill half the horizontal and vertical width of the screen `(960 × 540px)`, with a spatial frequency of `4cyc/960px` and temporal (drift) frequency of `1Hz`.

We observed a bimodal distribution of latencies (Quartiles: `28, 30, 36ms, n=50`, Figure 4.8), presumably because onsets of visual stimuli are quantized to the refresh rate `(60Hz, 16.67ms)` of the monitor. This range of latencies corresponds to the second and third frame after the trigger is sent (2/3 of observations fall in the 2nd frame, 1/3 of observations in the 3rd frame). We observed a median framerate of `36.2 FPS (IQR: 0.7)` across 50 trials (`8863` frames, Figure 4.9).

**Figure 4.8:**
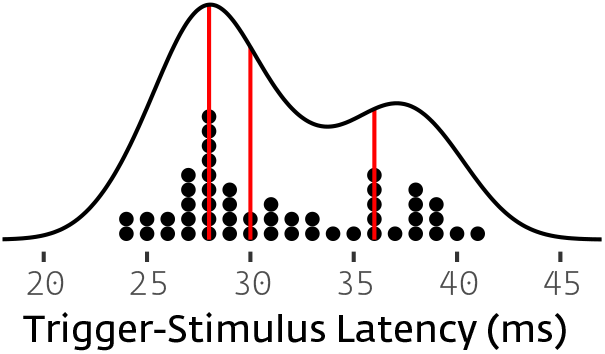
Stacked dots are a histogram of individual observations (n=50) underneath the probability density (black line), red lines indicate quartiles.

**Figure 4.9:**
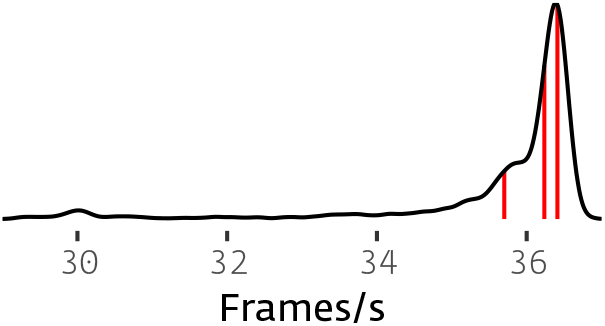
Probability density of framerates for 960 × 540px grating rendered at 1080p. Red lines indicate quartiles

We further tested the Pi’s framerate by using Psychopy’s [timeByFrames](https://github.com/psychopy/psychopy/blob/3.1/psychopy/demos/coder/timing/timeByFrames.py) test—a script that draws stimuli without any Autopilot components running—to see if the framerate limits were imposed by the hardware of the Raspberry Pi or overhead from Autopilot (Table 4.8). We tested a series of Gabor filters and [random dot stimuli](https://www.psychopy.org/api/visual/dotstim.html#psychopy.visual.DotStim) (dots travel in random directions with equal velocity, default parameters) at different screen resolutions and stimulus complexities. The Raspberry Pi was capable of moderately high framerates (`>60 FPS`) for smaller, lower resolution stimuli, but struggled (`<30 FPS`) for full HD, fullscreen stimuli.

**Table 4.8:**
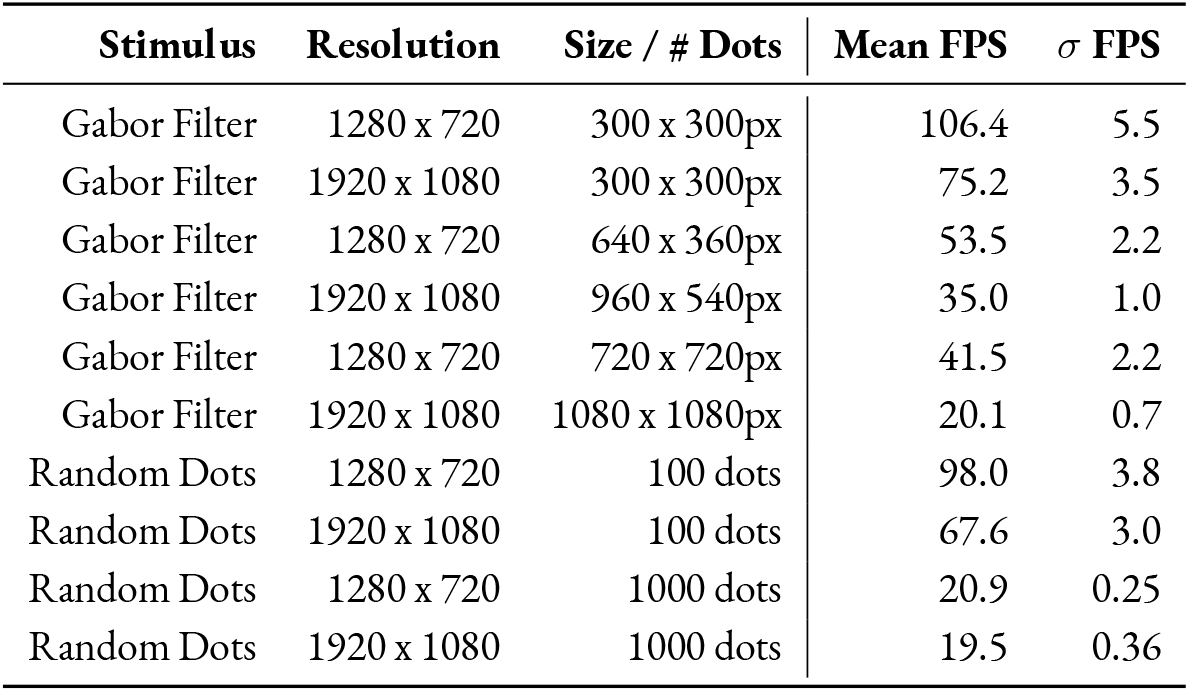
Tests performed over 1000 frames with PsychoPy’s [timeByFrames](https://github.com/psychopy/psychopy/blob/3.1/psychopy/demos/coder/timing/timeByFrames.py) test.

| Stimulus | Resolution | Size / # Dots | Mean FPS | $\sigma$ FPS |
| --- | --- | --- | --- | --- |
| Gabor Filter | 1280 x 720 | 300 x 300px | 106.4 | 5.5 |
| Gabor Filter | 1920 x 1080 | 300 x 300px | 75.2 | 3.5 |
| Gabor Filter | 1280 x 720 | 640 x 360px | 53.5 | 2.2 |
| Gabor Filter | 1920 x 1080 | 960 x 540px | 35.0 | 1.0 |
| Gabor Filter | 1280 x 720 | 720 x 720px | 41.5 | 2.2 |
| Gabor Filter | 1920 x 1080 | 1080 x 1080px | 20.1 | 0.7 |
| Random Dots | 1280 x 720 | 100 dots | 98.0 | 3.8 |
| Random Dots | 1920 x 1080 | 100 dots | 67.6 | 3.0 |
| Random Dots | 1280 x 720 | 1000 dots | 20.9 | 0.25 |
| Random Dots | 1920 x 1080 | 1000 dots | 19.5 | 0.36 |

Autopilot is appropriate for realtime rendering of simple stimuli, and the proof-of-concept API we built around Psychopy doesn’t impose discernible overhead (Mean framerate for a `960 × 540px` grating at `1080p` in Autopilot: `36.2 fps`, vs. `timeByFrames 35.0 fps`). In the future we will investigate prerendering and caching complex stimuli in order to increase performance. A straightforward option for higher-performance video would be to deploy an Autopilot agent running on a desktop computer with a high-performance GPU, or to use a single-board computer with a GPU like the [NVIDIA Jetson](https://www.nvidia.com/en-us/autonomous-machines/embedded-systems/jetson-nano/) ($99)^15^.

## 5 Limitations and Future Directions

We will likely never view Autopilot as “finished.” Autopilot—like all open-source software—is an evolving project, and this paper captures it as a snapshot at v0.5.0. We are invested in its development, and will be continually working to fix bugs, make its use more elegant, and add new features in collaboration with other researchers.

We expect that as the codebase matures and other researchers use Autopilot in new, unexpected ways that some fundamental elements of its structure may evolve. We have built version logging into the structure of the system so that changes will not compromise the replicability of experiments (see 5.5 below). While there will inevitably be breaking changes, these will be transparently documented, [announced in release notes](https://docs.auto-pi-lot.com/en/latest/changelog/index.html), and indicated with [semantic versioning](https://semver.org/) in order to alert users and describe how to adapt as needed.

We recognize the risk and inertia of retooling lab infrastructure, and there is still much work to be done on Autopilot. We welcome all [issues](https://github.com/auto-pi-lot/autopilot) and [questions](https://github.com/auto-pi-lot/autopilot/discussions) from anyone interested in contributing, or just curious to try it out — trying Autopilot is ultimately as risky as buying a Raspberry Pi.

The current major planned changes (also see the [todo](https://docs.auto-pi-lot.com/en/latest/todo.html) page in the docs) include:

1. **Python, Meet Rust** - Python is very useful as a high-level glue language, and its accessibility to a large number of scientific programmers is important to us, but it has its own very real performance limitations. As Autopilot’s modules mature and stabilize, we are interested in rewriting core routines like sound presentation and networking in rust and exposing them to python with tools like PyO3
2. **Real Realtime** - Beneath user space decisions like programming language, the timing of CPU operations in linux is still determined by the kernel — this is one of the major reasons why other projects are based around dedicated microcontrollers. For almost everything that most scientists want to do, the standard linux kernel is perfectly fine, but we are interested in investigating what it would take to provide true deterministic realtime performance via Autopilot’s high-level object system. One approach might be to provide [prebuilt realtime kernel](https://github.com/ros-realtime/linux-real-time-kernel-builder) images along with tools to easily deploy them, though no firm plan has been made.
3. **Integration** - We will continue to collaborate with other programming teams to be interoperable with a broader array of other tools. Our next set of planned integrations include recording electrophysiological data by integrating with Open Ephys[57], optical imaging data from the Miniscope project [58, 59], and shared processing and control pipelines with Bonsai[18].
4. **Data Ingest & Export** - We are releasing Autopilot’s data modeling system in v0.5.0 as an alpha release alongside this paper, and it includes prototype export interfaces to [Neurodata Without Borders](https://www.nwb.org/)[33] and [Datajoint](https://www.datajoint.com/)[60]. Over the next several releases, we will continue to improve our data model so that researchers can easily structure their data and choose among different backends for storage. We are also working on a [separate project](https://github.com/auto-pi-lot/ingesture) to make tools to ingest data from the more ad-hoc directory-based data formats widely used in science and ingest them into Autopilot’s and other tools formal modeling systems. In the longer term, we are interested in making Autopilot interoperable with linked data systems as part of a broader vision of digital infrastructure.
5. **Provenance** - Autopilot stores version information and local configuration in multiple places, and it is technically possible to faithfully replicate an experiment, but recording of provenance can still be consolidated and improved. By formalizing our object and data model, we will also systematize the many changes in configuration and version possible across the system for complete provenance tracking.
6. **P2P Networking** - The default tree structure of Autopilot’s networking modules has proven to be unnecessarily limiting over time. In part, we had preoptimized for processing messages in a separate processes assuming that would help problems from dropped messages and overflowing send buffers, but in practice messages are almost never dropped and network nodes are as effective as stations in sending and receiving large amounts of data. As part of unifying Autopilot’s object system, we will implement a fully peer-to-peer networking system such that each instantiated object has a unique ID so that messages can be easily addressed from any object to any other in its swarm. We will learn from previous p2p addressing systems like distributed hash tables to allow net nodes to join the swarm and discover all other nodes automatically without manually configuring IP addresses and ports. In the longer term we are interested in peer to peer data transfer as well, so that an object serving as a data source can efficiently stream to many consumers without needing to duplicate each message for every consumer.
7. **Slots, Signals, and Streaming** - We will be supplementing a more general network structure with a system of specifying which attributes of each object are data sources, which are sinks, and what kind of connection they accept. Similar to Qt’s [signal and slot](https://doc.qt.io/qt-6/signalsandslots.html) model, we want to make it as easy as using a `.connect()` method to control one piece of hardware with another. The transforms module should also be able to support branching and forking operations so multiple data sources can be combined for elaborated hardware control. ZeroMQ is an excellent tool for sending and receiving control messages, but formalized signals and slots could also specify different streaming tools like [redis](https://redis.io/) or [gstreamer](https://gstreamer.freedesktop.org/) that might be better suited for high-bandwidth linear streams like video. Applied generally, this could also solve related problems like the relatively implicit handling of event triggers in the Task class and the need for manual configuration of connections between pilots and copilots.
8. **Rebuild the GUI** - The GUI is some of the oldest code in the library, was written before most of the other modules existed, and needs to be rebuilt. We have started by remaking its central widgets to be [generated from pydantic models](https://github.com/auto-pi-lot/autopilot/blob/f0d20fbf3b33f832cf31136ab7d16abe01a3e924/autopilot/gui/widgets/model.py) used increasingly throughout the system, but the rest of the GUI still needs to be rearchitected into a structure that decreases code duplication and allows us to do things like provide GUI extensions via plugins. We will likely continue to use Qt for the near future, but are also exploring the idea of webassembly tools to make browser-based web interfaces for remote control.
9. **Plugins** - We want Autopilot’s plugin system to be permissive and as natural as the scripting style that most experimental code is written in, but we still need some means of specifying dependencies on other packages and plugins, among other improvements. We will be making a plugin generator that makes a folder of plugin boilerplate, as well as tools for installing, uploading, and synchronizing versions with git and the wiki. Over time we will make all object types within Autopilot able to be extended with plugins, as well as make it possible to override and extend built-in objects.
10. **Knowledge Organization** - We have been extending our thinking from code itself to more broadly consider the social systems that surround research code. The wiki was our first step, and we will continue to make more points of integration for smoothly incorporating contextual knowledge typically stored in lab notebooks into a public, collectively curated information system. We want to make it easier not only for individual researchers to use Autopilot, but make it easier for labs to coordinate work across projects without needing to rely on proprietary SaaS platforms with additional tooling for managing swarms, and moving beyond a single Autopilot wiki to a federated system of wikis for fluid continuity between “private” local coordination and “public” shared knowledge.
11. **Tests** - Our collection of tests doesn’t cover the whole codebase, and so as we formalize our contribution process will move towards a system where all new code must have tests and documentation to be integrated. We also want to integrate our tests more closely with our documentation so that researchers know which part of the code has explicit tests guaranteeing functionality.
12. **Security** - Autopilot is a networked program, and while it doesn’t execute arbitrary code from network messages, there is no security model to speak of. So far this hasn’t beren a problem, as we encourage only using Autopilot on a local network behind a router, but as we we build out our networking modules we will investigate how to incorporate identity verification systems to protect swarms from malicious messages.
13. **Metastructure & API Maturity** - The scope and structure of Autopilot is still in flux relative to other, more mature Python packages. To reach a stable v1.0.0 API, we are in the process of unifying Autopilot’s object structure so everything is clearly typed, all configuration is explicit, and all code written to handle special cases is absorbed into more general systems. Different parts of Autopilot have had different degrees of care over time, and so we will be working to catch the oldest modules up, trim unused ones, and make sure every line in the library is documented and useful. For the time being, flexibility is useful because frequently used or requested features trace a desire path outlining how its users believe Autopilot should behave. Each shortcoming we fix in Autopilot’s modules makes it more straightforward to fix the rest, and so once the major remaining work is completed we will transition to a more conservative pace of development that ensures the longevity of the project.

## 6 Glossary

**Agent** 3.7 The executable part of Autopilot. A set of startup routines (eg. opening a GUI or starting an audio server), runtime behavior (eg. opening as a window or running as a background system process), and event handling methods (ie. **listens**) that constitute the role of the particular Autopilot instance in the **swarm**.

**Copilot** 3.7 An **agent** that performs some auxiliary, supporting role in a **task—** primarily used for offloading some hardware responsibilities from a **pilot**.

**Graduation** 3.3 Moving between successive **tasks** in a **protocol** when some criterion is met.

**Listen** 3.8 A method belonging to the **station** or **node** of a particular **agent** that defines how to process a particular type of message (ie. a message with a particular key).

**Node** 3.8 A networking object that some module (eg. hardware, **tasks**, GUI routines) or method (eg. a **listen**) uses to communicate with other **nodes**. Messages to other **agents** in the swarm are relayed through their **Station**

**Pilot** 3.7 An **agent** that runs on a Raspberry Pi, the primary experimental agent of Autopilot. Typically runs as a system service, receives **tasks** from a **terminal** and runs them. Can organize a group of **children** if requested by the **task**.

**Protocol** 3.3 A (.json) file that contains a list of **task** parameters and the **graduation** criteria to move between them. The **tasks** in a protocol are also known as its **levels**.

**Stage** 3.3 **Stages** are methods that implement the logic of a **task**. They can be used analogously to states in a finite-state machine (eg. wait for **trial** initiation, play stimulus, etc.) or asynchronously (whenever x input is received, rotate stimulus by y degrees).

**Station** 3.8 Each **agent** has a single **station**, a networking object that is run in its own process and is responsible for communication between **agents**. The **station** also routes messages from **children** or other **nodes**.

**Swarm** Informally, a group of connected **agents**.

**Task** 3.3 A formalized description of an experiment: the parameters it takes, the data that it collects, the hardware it needs, and a collection of **stages** that describe what happens during the experiment.

**Terminal** 3.7 A user-facing **agent** that provides a GUI for operating and maintaining a **swarm**.

**Topology** 3.7 A particular combination of **agents**, their designated responsibilities, and the networking connections between them invoked by a **task** (eg. task requires one pilot to record video, one to process the video, and one to administer reward) or by usage (eg. 10 pilots are connected to a single terminal and are typically used to run 10 independent tasks, though they could run shared tasks together).

**Trial** 3.3 If a **task** is structured such that its **stages** form a repeating series, a **trial** is a single completion of that series.

## Acknowledgements

*We would like to acknowledge and thank Lucas Ott and Tillie Morris for doing most of the behavioral training and being so patient with the bugs, Brynna Paros and Nick Sattler for their help with constructing our behavioral boxes, Chris Rogers who has been brave enough to adopt and contribute to Autopilot in its roughest state, Arne Meyer, Mikkel Roald-Arbøl, and David Robbe who have contributed code and advice, Mackenzie Mathis, Alex Mathis, Gonçalo Lopes, and Gary Kane who collaborated on the DeepLabCut-Live project and provided many a mentorship along the way, Jeremy Delahanty for his inspiring tenacity and humbleness in thinking about better research tools, Matt Smear and Reese Findley for loaning us their Bpod for far longer than they intended to, John Boosinger and the rest of the staff in the machine shop for all their advice and letting me use all their tools, Erik Flister whose Ratrix software inspired some of the design features of Autopilot [1], my labmates Molly Shallow and Sam Mehan who kept me afloat in my last months of dissertation writing, Rocky Penick for her help strapping Autopilot onto the back of Evan Vickers’ mesoscope rig (and Evan for letting us play with his rig), several artists on* flaticon.com *([Freepik](https://www.flaticon.com/authors/freepik), [Nikita Golubev](https://www.flaticon.com/authors/nikita-golubev), [Those Icons](https://www.flaticon.com/authors/those-icons)) whose work served as stems for some of the figures, and the Janet Smith House for the endless support and relentless criticism of the figures. This material is based on work supported by NIH NIDCD R01 DC-015828, NSF Graduate Research Fellowship No. 1309047, and a University of Oregon Incubating Interdisciplinary Initiatives award*.

## Contribution Statement

JLS designed and wrote the software, documentation, wiki, figures, ran the tests, and wrote and edited the paper. LO trained the animals and did an extensive amount of beta testing, bugfinding, and made some of the hardware designs on the wiki. MW mentored, edited the paper, and beta tested the software.

## Footnotes

1 “Our original definition of groupware was ‘intentional group processes plus software to support them.’ It has both *computer* and *human* components: software of the computer and ‘software’of the people using it. […] Recently this definition has been extended to include other more expressly cultural factors including myth, values and norms. The computer software should reflect and support a group’s purpose, process and culture.” Peter and Trudy Johnson-Lenz (1991)[14]

2 See Table 3.2

3 https://docs.auto-pi-lot.com

4 https://wiki.auto-pi-lot.com

5 **Other tools:** - Bonsai[18] - [site](https://bonsai-rx.org/), [git](https://github.com/bonsai-rx/bonsai) - Expyriment[19] - [site](https://www.expyriment.org/), [git](https://github.com/expyriment/expyriment.git) - PsychoPy[20] - [site](https://www.psychopy.org/), [git](https://github.com/psychopy/psychopy) - OpenSesame[21] - [site](https://osdoc.cogsci.nl/), [git](https://github.com/smathot/OpenSesame) - SMiLE - [docs](https://smile-docs.readthedocs.io/en/latest/) - ArControl[22] - [git](https://github.com/chenxinfeng4/ArControl) - and see [OpenBehavior](http://openbehavior.com/)

6 Bpod runs [custom firmware](https://github.com/sanworks/Bpod_StateMachine_Firmware) written in C++ on a [Teensy 3.6](https://www.pjrc.com/store/teensy36.html) microcontroller. pyControl’s pyboard runs [micropython](https://micropython.org/), a subset of Python that excludes canonical libraries like numpy[23] or scipy[24]

7 And Autopilot, of course, also has many of its own weaknesses

1 [Raspberry Pi model 4B](https://www.raspberrypi.org/products/raspberry-pi-4-model-b/), see Table 3.2

2 and improvements to CPython in Python 3.11 and onwards will bring overhead close to zero[28]

3 See David Beazley’s ‘[Understanding the Global Interpreter Lock](http://www.dabeaz.com/python/UnderstandingGIL.pdf)’ and associated [visualizations](http://www.dabeaz.com/GIL/gilvis/index.html).

4 The user guide and API documentation are available at docs.auto-pi-lot.com

5 For readability of the docs, we omit generating HTML documentation for some private methods and functions, but they are documented in the source and their function is made clear from their context and the documentation of public methods.

6 Like inheriting from the [GPIO](https://docs.auto-pi-lot.com/en/latest/hardware/gpio.html#autopilot.hardware.gpio.GPIO) class gives GPIO plugins a systematic means of interacting with the underlying pigpiod daemon.

7 “ZeroMQ […] has a subversive effect on how you develop network-capable applications. […] message processing rapidly becomes the central loop, and your application soon breaks down into a set of message processing tasks.” “If there’s one lesson we’ve learned from 30+ years of concurrent programming, it is: *just don’t share state*.” > -[The ZeroMQ Guide](http://zguide.zeromq.org/)

8 Though our Subject class provides a simplified interface to access and manipulate Autopilot data

9 Coverage statistics for Autopilot are available on coveralls.io at https://coveralls.io/github/auto-pi-lot/autopilot

1 with care for backwards compatibility

2 Released as an alpha version at the time of writing

3 An example of subclassing a generic ‘Task’ class is included in Autopilot’s [user guide](https://docs.auto-pi-lot.com/en/latest/examples.html)

4 We take inspiration from Aaron Swartz’ description of another engineering project, the Semantic Web, that became too precious about its formalisms: “Instead of the “let’s just build something that works” attitude that made the Web (and the Internet) such a roaring success […] they formed committees to form working groups to write drafts of ontologies that carefully listed (in 100-page Word documents) all possible things in the universe and the various properties they could have, and they spent hours in Talmudic debates over whether a washing machine was a kitchen appliance or a household cleaning device. […] And instead of spending time building things, they’ve convinced people interested in these ideas that the first thing we need to do is write *standards*. (To engineers, this is absurd from the start—standards are things you write *after* you’ve got something working, not before!)”[48]

5 a thin wrapper around scipy’s signal.gammatone

6 though the Raspberry Pi compute module has a PCI lane that supports GPUs.

7 As of v0.5.0, Autopilot is packaged with [Poetry](https://python-poetry.org/), so they are `[tool.poetry.extras]` entries within the `pyproject.toml` file, installed with pip like `pip install auto-pi-lot[pilot]` or poetry like `poetry install -E pilot`

8 A previous version of this paper described a third, subordinate “Child” agent that performed auxiliary operations in a task. We now view such a hierarchy as unnecessary, and that distribution of labor within a task is better served by a fluid combination of multiple Pilots than thinking of them as qualitatively different agents. We now refer one among multiple agents performing a task together as a “copilot.”

9 Autopilot uses ZeroMQ[31] and [tornado](http://www.tornadoweb.org/en/stable/) to send and process messages

10 Though automatically configuring the use of faster protocols like IPC for communication within an agent or different backends like redis or gstreamer for data streams that would benefit from them is part of our development goals

11 converted to binary suitable for sending between computers

12 For example, to send a message from E to C in the diagram above: 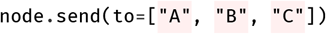

13 Autopilot uses [PySide](https://wiki.qt.io/PySide), a wrapper around [Qt](https://www.qt.io/), to build its GUI.

1 All procedures were performed in accordance with National Institutes of Health guidelines, as approved by the University of Oregon Institutional Animal Care and Use Committee.

2 A previous version of this paper included benchmarking and comparison to Bpod and pyControl’s sound onset latency, but since then both packages have changed substantially, including Bpod creating a new [hifi sound module](https://sites.google.com/site/bpoddocumentation/assembling-bpod/hifi-module) based off HiFiBerry hardware very similar to the card used here, making those benchmarks obsolete. In this version we have omitted comparative benchmarks in favor of allowing the maintainers of those packages to publish their own benchmarks.

3 Our sync is likely to be near to or better than that reported in [56]: in addition to a quiet network, we configured chrony to poll more frequently and tolerate a smaller error than default

4 (1920 * 1080*3*8 bits) / 8 = 6-megabytes per frame of a 1080p color video, which is why video is rarely streamed uncompressed

5 maximum average rate of 5000 messages for each of the equivalent empty message tests in the four conditions described below

6 In all cases, float64 numpy arrays encoded in base64

7 n=5,000 for each condition at each size

8 A message with a 1-MByte zero array payload compressed to 6-KBytes.

9 In this dataset. There is additional payload bandwidth headroom with larger messages, and we include an additional dataset with a 200MByte/s bandwidth in the supplement.

10 2-Photon: 5.9MB/s (12 bits * 512×512 resolution* 15Hz)

11 Neuropixels: 14.4MB/s[6] (10 bits * 30kHz * 384 channels)

12 No mice were trained on this task

13 `(1920×1080px, 60Hz)`

14 Visual stimuli were presented with [Psychopy](https://www.psychopy.org/) using the [glfw](https://www.glfw.org/) backend while Autopilot was run in a dedicated X11 server.

15 as we did in [30]

## Notes

### Competing Interest Statement

The authors have declared no competing interest.

### Summary of Updates

Updates to software design and implementation

